# Heatwaves rescue a mosquito host from parasitism across a large geographic gradient

**DOI:** 10.64898/2026.06.26.734620

**Authors:** Johannah E. Farner, Izzy M. Riley, Aspen H. Singh, Erin A. Mordecai

**Affiliations:** Biology Department, Stanford University, Stanford, CA, USA

**Keywords:** parasitism, temperature, heatwave, climate change, local adaptation, *Aedes sierrensis*, *Lambornella clarki*, mosquito, ciliate

## Abstract

The impacts of increasingly frequent and intense heatwaves on parasitism are an important frontier for understanding disease risk under climate change. These impacts are complex because parasitism arises from multiple interacting host and parasite traits that can vary in thermal sensitivity, across the ontogeny of infection, and among populations exposed to different historical temperature regimes. We used a lab microcosm experiment to investigate the effects of heatwaves occurring during two different phases of a mosquito host – ciliate parasite interaction that occurs during the winter in western North America, for six host – parasite populations from two geographic regions with differing histories of winter heat. We found that because heatwaves allowed hosts to evade infection, they reduced parasitism and increased survival. An early heatwave during initial parasite attack had stronger effects than a later heatwave occurring after infection establishment. We did not find evidence of locally adapted heatwave effects on parasitism. However, heatwave impacts were mechanistically predictable from previously measured thermal performance curves that described lower infection and stronger host defenses at warm constant temperatures. The results suggest that heatwaves may accelerate geographic shifts in parasitism, and demonstrate how fundamental host – parasite thermal biology links to the impacts of extreme temperature events.

## Introduction

Host – parasite interactions are widespread in nature, biologically important, and temperature-sensitive, making the effects of rapid global temperature change on disease a key research area in ecology (Altizer et al. 2013; Rohr & Cohen 2020). In addition to warming average temperatures, the frequencies and intensities of extreme temperature events such as heatwaves and cold snaps are increasing (Seneviratne et al. 2021). Both gradual warming and heat anomalies can alter the risk and severity of disease outbreaks (Cohen et al. 2020; Karvonen et al. 2010; Pounds et al. 2006). However, most research has focused on constant and mean temperatures, while understanding of how short-term temperature extremes impact disease risk remains comparatively limited (Claar & Wood 2020; Rohr & Cohen 2020). Because weather events such as heatwaves are transient and host – parasite interactions are dynamic, impacts on infection may depend on how temperature variability intersects with both the nonlinear thermal responses of host and parasite traits and the ontogeny of infection (Altman *et al*. 2016). Addressing this knowledge gap is critical to anticipating both immediate and long-term effects of climate change on disease risk (Buckley et al. 2023; Kingsolver et al. 2015; Kingsolver & Buckley 2017; Molnár et al. 2013; Mordecai et al. 2019).

The challenge of predicting heatwave impacts on infection is illustrated by the complex findings of studies examining parasitism under variable temperatures. All possible effects of heatwaves on parasitism have been observed, with no clear pattern among systems or experimental conditions where infections increased (Eisenlord et al. 2016; Green et al. 2019; Studer & Poulin 2013; Taig et al. 2025), decreased (Greenspan et al. 2017; Morales-Serna et al. 2021; Shodipo et al. 2020), or remained constant (Kunze et al. 2022; Landis et al. 2012; Pounds et al. 2006; Simaz & Szűcs 2021). Moreover, outcomes following heatwaves have been shown to differ within study systems depending on experimental factors such as heatwave amplitude, duration, and timing; background temperature conditions; and the experimental host and parasite genotypes or populations used (Dallas & Drake 2016; Kunze et al. 2022; Malinski et al. 2025; Manlik et al. 2023; McCartan et al. 2025, 2026; Rozmann et al. 2026).

Understanding complex heatwave effects on parasitism may require integrating underlying trait thermal responses and the natural history of host – parasite interactions with ecologically relevant heatwave treatments across populations. Lab-measured thermal performance curves (TPCs) that describe the frequently nonlinear and distinct thermal responses of hosts, parasites, and their interactions to constant temperatures may be particularly informative (Cohen et al. 2017; Rohr & Cohen 2020). Whether heatwaves shift conditions towards or away from thermal optima for key host and parasite traits may predict when hosts, parasites, or neither will be favored. For example, heatwaves might benefit hosts by boosting immune defenses or reducing parasite survival; benefit parasites by increasing metabolic demands on hosts or the speed of parasite attack or reproduction; or benefit neither if species-specific benefits or harms of heat exposure are offsetting, shared, or similarly harmful for host and parasite (Claar & Wood 2020; Ragonese et al. 2024). The timing of heatwaves in the development and infection cycles of hosts and parasites may also alter their overall impact as the relative importances and thermal responses of different traits shift, and as the influence of pre-heatwave conditions accumulates (Altman et al. 2016; Li et al. 2025; Raffel et al. 2013). Trait acclimation to temperature spikes may also change throughout the ontogeny of infection (Li et al. 2025; Pottier et al. 2026). Finally, effects may vary among populations adapted to different historical climates and heatwave regimes. However, linkages between heatwave effects and TPCs throughout the course of disease progression and among populations from different climates remain untested.

This study addresses the question of whether TPCs that describe relationships between parasitism and constant temperatures can mechanistically predict complex heatwave impacts for populations from across a climatically diverse species range. We use an ectothermic, winter-adapted system of the western tree hole mosquito host, *Aedes sierrensis*, and its facultative ciliate parasite, *Lambornella clarki*, that is widespread across a large geographic area on the west coast of North America; has a multi-stage infection process that can be tracked in lab experiments; and for which we have previously characterized the distinct, nonlinear TPCs of the host - parasite interaction and underlying traits for populations from across the species range (Couper et al. 2024; Darsie & Ward 2005; Farner et al. 2026; Ismail et al. 2023; Lyberger et al. 2024 Washburn *et al*. 1988a; Washburn & Anderson 1986) (Figure 1A, B). These features of the *Ae. sierrensis* – *L. clarki* system present a unique opportunity to develop and test hypotheses of how heatwaves and their timing might affect parasitism, given host and parasite thermal biology and the seasonal progression of ciliate and mosquito activity within tree holes.

**Figure 1.**
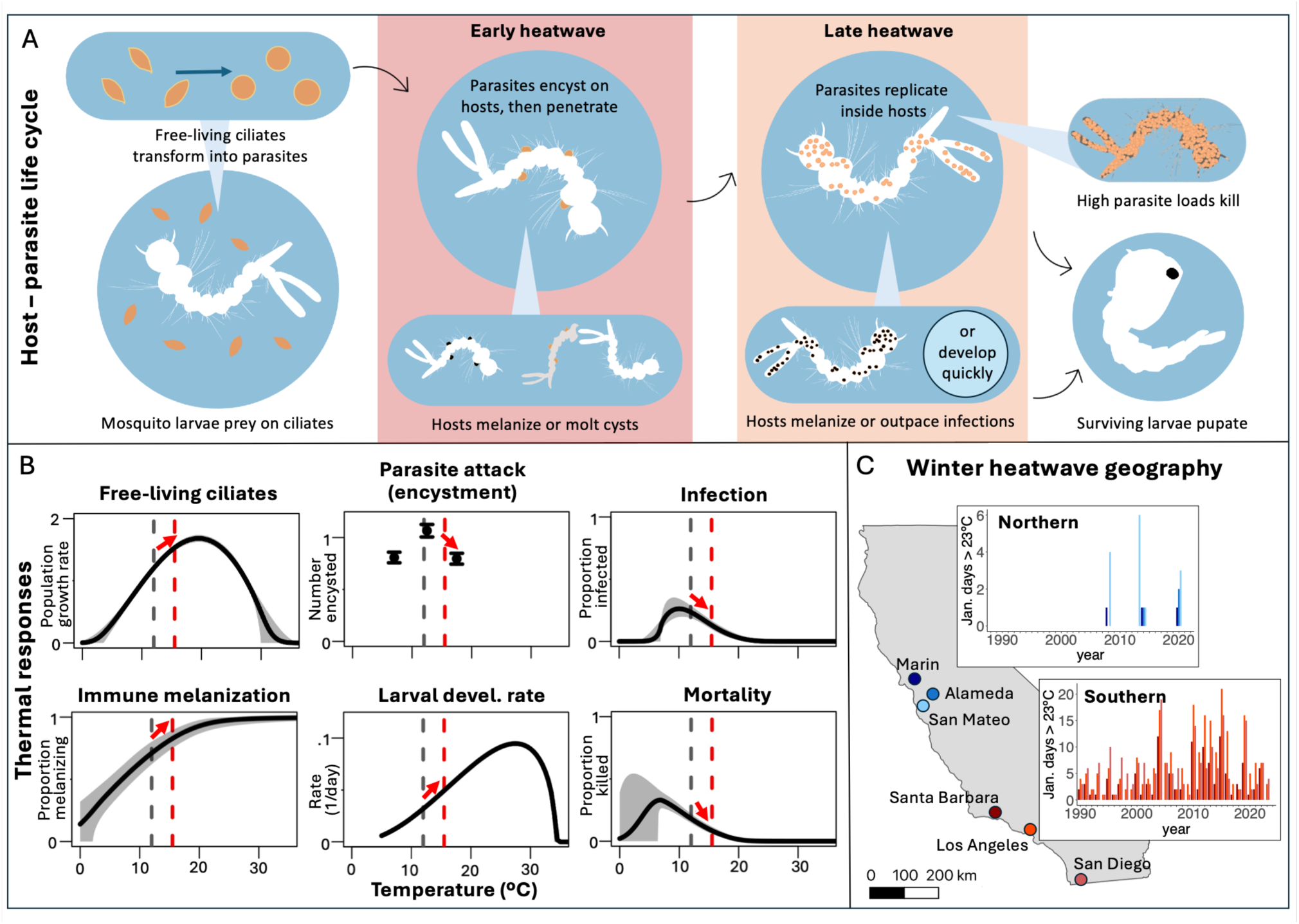
Heatwave impacts on parasitism may depend on how heat timing intersects with host – parasite life cycles, thermal responses, and population histories of heat exposure. (A) The interaction between the host mosquito *Aedes sierrensis* and facultative ciliate parasite *Lambornella clarki* progresses through distinct phases of parasite attack and infection (top row), which may respond differently to discrete heatwaves (colored boxes). Host defenses include an active melanization immune response and passive accelerated development at warm temperatures (bottom row). Mosquitoes surviving the host – parasite interaction pupate before emerging as adults. (B) Red arrows indicate hypothesized effects of heatwaves on parasite traits, host defenses, and interaction outcomes based on previously measured responses to constant temperatures (black curves and points). Gray dashed lines show the mean baseline temperature, and red dashed lines show the mean temperature during heatwave treatments in the experiment. Thermal response data are replotted from Couper et al. (2024), Lyberger et al. (2024), and Farner et al. (2026). (C) Six experimental host – parasite populations were sourced from two climatic regions with differing historical winter heatwave frequencies. Blue shades show northern and red shades show southern field sites in California, USA. Bar charts show counts of January days with maximum temperatures exceeding 23°C from 1988 to 2023 recorded by the weather station nearest to each field site with available data.

In previous field surveys and lab experiments, we have found that infection is most likely to occur at cool mean and constant temperatures where host defenses are suppressed (Farner et al. 2026; Ismail et al. 2023; Lyberger et al. 2024). Thus, we hypothesized that (H1) heatwaves would decrease infection rates and increase host survival, by strengthening host immune and developmental defenses (Figure 1B). However, the host – parasite interaction progresses through distinct stages of parasite attack, infection, and replication, such that stronger host defenses may be required to clear infections at later stages. Larval mosquitoes can escape attacking parasites through a strong initial melanization immune response, or if they develop quickly (Figure 1A) (see Methods for details). However, once infections are established, heatwaves might benefit both host defenses and parasite replication within hosts. Therefore, we also hypothesized that (H2) heatwaves would have stronger effects when imposed during parasite attack compared to after infections had established. Finally, local adaptation to heatwaves might mediate their impacts. We have found that although the lab-measured 10°C constant temperature thermal optimum for infection is not locally adapted, parasitism in warm climates that regularly exceed this temperature is reduced but persistent (Farner et al. 2026; Lyberger et al. 2023). Additionally, thermal optima for free-living *L. clarki* ciliates correlate with local mean temperatures (Ismail et al. 2023; Lyberger et al. 2024). Based on these indications of potential parasite thermal adaptation, we hypothesized that (H3) heatwave impacts would be stronger for populations from climates where warm winter days are rare (Farner et al. 2026; Ismail et al. 2023; Lyberger et al. 2024) (Figure 1C). To test these hypotheses, we used a laboratory experiment with ecologically relevant heatwave treatments applied either early or late in the host – parasite interaction, for six *Ae. sierrensis* and *L. clarki* host – parasite populations from two different climatic regions corresponding to either rare or common historical heat exposure.

## Methods

### Study system

*Aedes sierrensis* and *L. clarki* occur across western North America in tree hole habitats, commonly in oak woodlands, oak savannas, and riparian habitats (Darsie & Ward 2005; Washburn & Anderson 1986). The host – parasite interaction follows a seasonal progression that initiates when early winter rains fill tree holes and aquatic mosquito larvae and free-living ciliates hatch (Darsie & Ward 2005; Washburn & Anderson 1986) (Figure 1A). The mosquito larvae prey on ciliates present in the water column, and free-living ciliates facultatively transform into a parasitic morph that attacks the larvae within a few days of tree hole filling (Washburn et al. 1988b). Infections occur when parasitic *L. clarki* encyst on the mosquito cuticle and then successfully enter the body cavity, where the ciliates replicate, often generating lethal densities of daughter cells within a few weeks (Washburn et al. 1988a). Hosts directly defend against parasites with a melanization immune response, and indirectly through accelerated development at warm temperatures (Corliss & Coats 1976; Farner et al. 2026; Washburn et al. 1988b, 1991b). Melanization is a generalized insect immune response that encapsulates invading cells in melanin (Christensen *et al*. 2005). When successful, melanization of attacking parasites forming cuticular cysts can prevent infection. Later on, melanization of infecting parasites within the host body cavity can clear infections (Corliss & Coats 1976). Melanization occurrence and strength varies among populations and increases with temperature (Farner et al. 2026, Lyberger et al. 2024). At warm temperatures, accelerated host development also acts as a passive defense mechanism. As hosts progress through four larval instars before pupating, shorter intervals between molts can increase the likelihood that ciliates attached to the cuticle are shed before penetrating into the body cavity (Washburn et al. 1991b). Faster host development also limits the time window for internal infections to intensify and kill larvae before pupation (Farner et al. 2026). Mosquitoes undergo one developmental cycle per year, with adults emerging and laying eggs in the spring (Hawley 1985). Free-living ciliates are released back into tree holes when infected larvae die, and can disperse to new tree holes via infected mosquitoes that survive to adulthood (Egerter et al. 1986).

We have previously used field surveys and TPC experiments to characterize the thermal biology of this system. This work has shown that parasitism is broadly adapted to average winter temperatures across much of the species range, with infections most likely to occur at cool temperatures that suppress host defenses. Infection responds unimodally to temperature, peaking at 10°C average wet season temperatures in field observations and at 10°C constant incubation temperatures in laboratory experiments (Farner et al. 2026) (Fig. 1B). The 10°C thermal optimum for parasitism is cool compared to the optimal temperatures for the mosquito’s defenses: melanization increases with temperature up to a plateau near 22°C, and the larval development rate peaks at 27.2°C (Couper et al. 2024; Farner et al. 2026; Lyberger et al. 2024) (Fig. 1B). Thermal optima for free-living *L. clarki* population growth are also comparatively warm (Figure 1B) and exhibit local thermal adaptation (16°C – 24.5°C, depending on the parasite population) (Farner et al. 2026; Ismail et al. 2023; Lyberger et al. 2024).

### Materials

The mosquitoes and ciliates used in the July 2024 lab experiment originated from tree holes sampled in January and February 2024 at six field sites in two different climatic regions. Three sites were located in southern California (in San Diego, Los Angeles, and Santa Barbara counties), where hot winter days with temperatures exceeding 23°C are common, and three were located in northern California (in San Mateo, Alameda, and Marin counties), where winter heatwaves are comparatively rare, but increasing (Figure 1C; Tables S1, S2). Over two years of previous field surveys, parasite infections were more common in the northern region (21 – 36% of tree holes) compared to the southern region (11-16% of tree holes) (Farner et al. 2026). Each site was at least 40 km from all other sites. The average flight distance of *Aedes* mosquitoes is 89m, making substantial gene flow between the source locations of the experimental populations unlikely (Verdonschot & Besse-Lototskaya 2014).

Tree hole water samples containing mosquito larvae (N = 1 – 400 individuals per tree hole) were transported to Stanford University and inspected for *L. clarki* infection at 40x magnification. Larvae from tree holes where infections were absent were reared to adulthood and their eggs were used in the experiment. In parallel, ciliates were cultured from infected mosquitoes collected from different tree holes in the same sites. All experiments were conducted with sympatric mosquitoes and ciliates reared from field-collected samples. For each site, field-collected mosquitoes from three different uninfected tree holes containing at least 100 larvae each were reared together to ensure representative genetic diversity and sufficient egg production for the experiment. Ciliate cultures from two tree holes at each site were started from individual infected mosquitoes and maintained separately until the start of the experiment, then combined in a 1:1 ratio to create an inoculum. Field-collected mosquitoes and ciliates were maintained at 20°C in laboratory conditions, and mosquito eggs were matured for at least one month at 4°C. Mosquito rearing and parasite culturing followed the protocols in Ismail et al. (2023), Couper et al. (2024), and Lyberger et al. (2024).

### Temperature treatments

We chose diurnally fluctuating baseline and heatwave temperature conditions based on analysis of 35 years (1988 - 2023) of daily high and low temperatures from PRISM 4-km resolution data (PRISM Climate Group 2021). We have previously shown that PRISM temperature estimates closely track actual temperatures logged by iButtons placed inside tree hole habitats across the study area, which averaged 1°C cooler (Farner et al. 2026). In selecting experimental incubation temperatures, we focused on January temperatures to match the seasonal onset of *Ae. sierrensis* and *L. clarki* activity in tree holes in the study area. The baseline daily temperature fluctuation consisted of 12 daytime hours at 17°C and 12 nighttime hours at 7°C, and was based on average January daily maximum and minimum temperatures across all population source locations (Figure S1). In the heatwave treatments, the daytime temperature was raised to 24°C for two consecutive days. January temperatures had reached 24°C at all field sites at least once in the 35 years prior to the experiment, and were historically rare in the northern region (99^th^ – 100^th^ percentile of January daily high temperatures) compared to the southern region (83^rd^ – 88^th^ percentile). The heatwave nighttime temperature was kept at 7°C to match the 7.4°C average daily minimum temperature associated with hot January days (reaching 22°C to 26°C) across the field sites. In addition to reflecting natural temperature fluctuations during winter heatwaves, we expected the cool nighttime heatwave temperature to produce conservative estimates of heatwave impacts by providing a recovery period for heat-exposed hosts and parasites. A detailed explanation of the temperature selection procedure is provided in the Supplement.

### Heatwave experiment

We submerged 800 mosquito eggs laid by field-collected adults for each of the six study populations at room temperature (20°C), following the *Ae. sierrensis* hatching protocol in Couper et al. (2024). Groups of five larvae ≤48 hours old were sorted into wells in six-well plates (Corning #3736) and then placed in incubators. Each well contained one autoclaved barley seed and either 5 mL sympatric *L. clarki* parasite culture diluted to 35 cells per 100 µL plus 2 mL autoclaved culturing media (infection treatment), or 7 mL autoclaved culturing media (parasite control treatment). Wells with and without parasites were randomly assigned within each well plate. Well plates were assigned to receive either an early heatwave treatment, a late heatwave treatment, or a temperature control treatment with no heatwave (for six total heatwave x parasite exposure treatments). The early heatwave treatment, imposed seven days into the experiment, targeted the initial parasite attack phase, when *L. clarki* attaches to and attempts to penetrate the cuticle of early instar mosquito larvae. In contrast, the late heatwave treatment, imposed 21 days into the experiment, targeted established internal infections with parasite replication occurring inside the body cavities of late instar hosts.

Beginning four days into the experiment, we checked the mosquitoes weekly for survival and pupation, and used a dissection microscope (10 – 40X magnification) to observe (1) the proportion of larvae with cuticular ciliate cysts, corresponding to the first stage of *L. clarki* attack, when hosts may still evade infections through melanization and/or molting; (2) the proportion of larvae with established internal parasite infections, when host recovery requires that the immune melanization response and/or larval development to pupation outpaces parasite replication; and (3) the proportion of larvae with melanization spots present, indicating host immune recognition of attacking and/or infecting parasites (Washburn et al. 1988b, 1991b). Throughout the experiment, new observations of melanization represented hosts newly defending against attacking parasites or internal infections, and reduced fractions of melanizing hosts were caused by larval molting of cuticles with melanized cysts, infection clearance, and pupation. At each checkpoint, dead larvae were removed from wells and inspected for infection. Infections in dead larvae were included in the analyzed infection counts for the checkpoint at which they were observed, plus all subsequent checkpoints.

In all, 186 and 130 wells containing five mosquito larvae each were incubated with sympatric *L. clarki* present and absent, respectively. Across the experimental populations, the final dataset included 3 – 14 wells per combination of population, parasite exposure, and heatwave treatment (mean wells = 7.8, SD = 2.5) (Table S3). Differences in sample sizes resulted from both variation in initial mosquito egg hatching rates and sample loss due to human error during the experiment. The experiment ended after 12 weeks, when no surviving larvae remained. All pupation occurred within the first nine weeks.

Two populations had very low infection occurrence (<10 total at any point in the experiment) across all treatments: the northern population from Marin County, and the southern population from Los Angeles County. Both of these were excluded from all analyses presented in the main text, except where noted below. To explore host adaptation to local parasites as a potential source of low infection occurrence, we ran a supplemental experiment crossing allopatric host and parasite populations, described in the Supplemental Information.

### Statistical analyses

To assess how the early and late heatwave treatments affected *L. clarki* parasitism on *Ae. sierrensis* hosts, whether these effects translated into differences in host survival, and whether heatwave impacts varied between geographic regions with different historical heatwave regimes, we used generalized linear mixed effects models. Our main analyses focused on the hypothesized differences in heatwave impacts between geographic regions (northern versus southern) and between different phases of the host – parasite interaction (early versus late heatwaves compared to controls). Population fixed effects were not included in these models because replication was insufficient for heatwave impacts to be tested at both geographic scales simultaneously. Therefore, we checked the robustness of the results to potential subregional variation by fitting models that tested specifically for differences among populations (see Supplement: Methods for details). The results of these robustness checks were largely consistent with those of the main models; any differences are reported in the Results.

### Parasite attack (encystment)

We first investigated whether parasite attack (encystment) over the first two checkpoints responded to the early heatwave and/or differed between regions in ways that may have shaped subsequent larval infection levels. For this analysis, we modeled the proportion of live larvae with visible cuticular cysts as a beta-binomial distribution to account for overdispersion (Bolker 2008). Heatwave treatment, checkpoint, geographic region, and their interactions were included as fixed effects, and wells nested within populations were included as random effects. The late heatwave and temperature control treatments were grouped together for comparison with the early heatwave treatment because the late heatwave occurred after the relevant parasite attack period.

A supplemental analysis tested whether the populations excluded from the main model due to very low infection occurrence also experienced low parasite attack. This model included the data from all six populations, tested for interactions between population, heatwave treatment, and checkpoint, and included well ID as a random effect.

### Infection

We next tested for effects of the categorical variables—heatwave treatment, checkpoint, and geographic region—on the proportion of parasite-exposed larvae with active or fatal internal infections, out of the starting number of larvae. This analysis began at the second weekly checkpoint, when infections were first observed in a majority of populations, and ended at the fifth checkpoint, when a majority of larvae had either pupated or died (Figure S2). Checkpoint was included as a categorical variable because we expected the direction of its effect to differ before and after heat exposure. Later checkpoints were excluded because subsequent changes in larval infection were minimal, and increased model complexity with additional checkpoints was exacerbated by data loss as pupation continued. Wells nested within populations were included as random effects.

### Host immune defense

Next, we investigated whether the experimental heatwaves led to increases in the proportion of live larvae exhibiting a melanization immune response to *L. clarki* parasites. The host melanization models followed the structure of the three infection models described above, but the population random effects were dropped from the final model because these explained very little variance and prevented model convergence (Barr et al. 2013).

### Host development speed

Although not a direct response to parasites, faster larval development at warmer temperatures can also protect hosts from infection occurrence or lethality (Farner et al. 2026; Washburn et al. 1991b). To assess how the larval development rate was affected by parasite exposure, heatwave treatment, geographic region, and their interactions, we used discrete time-to-event analysis. Here, we modelled pupation hazard during the relevant time window (weeks 4 – 9) as a binomial distribution with a complementary log-log link (Austin 2017). Most pupation occurred within the first half of this period (98%). To enable inclusion of developmentally lagged hosts in the analysis, data from weeks seven (13 pupae), eight (3 pupae), and nine (1 pupa) were combined. To assess the hypothesis that parasite exposure, heatwave treatment, and geographic region might affect larval development rates independently and in combination, these predictors and all possible interactions were included in the model as fixed effects. To examine whether pupation hazard varied within the five week time window analyzed, checkpoint (week 4, 5, 6, or 7+) was included as an additive effect, with the fourth week treated as the reference level. Wells nested within populations were included as random effects. Because pupation hazard varied by checkpoint, we also examined its two-way interactions with parasite exposure, heatwave treatment, and geographic region. The final model included checkpoint x parasite exposure and checkpoint x heatwave treatment interactions that improved the model fit, as indicated by AIC values and likelihood ratio tests (χ² = 31.15, Bonferroni-adjusted p-value < 0.001).

### Host survival

Finally, we assessed whether host survival differed when mosquitoes were exposed to parasites and/or heatwaves, and whether these responses varied by geographic region. This model used the total proportion of mosquitoes surviving to pupation as the dependent variable and followed the formulation of the infection, encystment, and melanization models, adjusted to include parasite exposure and its interactions with other predictors as fixed effects, and to reflect the single observation per well.

### Model specification and post-hoc tests

For all models, the heatwave treatment predictor was encoded using treatment contrasts with the control treatment as the reference level, and sum-to-zero contrasts, which test for significant deviations from the overall mean, were used for all other categorical variables except where otherwise noted (Bolker 2018; Schielzeth 2010). Where possible, binomial distributions were fit with the R package “lme4” (Bates et al. 2003). As noted above, alternative distributions were fit as needed with the R package “glmmTMB” (Brooks et al. 2017). We used the R package “DHARMa” to perform model diagnostics and test for overdispersion (Hartig 2017). To follow up on significant interactions, we used pairwise comparisons with Bonferroni-adjusted p-values in the R package “emmeans” (Lenth 2023).

We used R version 4.6.1 for all analyses (R Core Team, 2022). In addition to the R packages cited above, we used “broom.mixed”, “contrastable”, “tidyverse”, “blme”, and “arm” for analysis, visualization, and formatting of data and results (Bolker et al. 2019; Dorie and Dorie 2015; Gelman et al. 2015; Sostarics 2024; Wickham et al. 2019).

## Results

### Parasite attack

Parasites quickly formed cysts on host cuticles at the start of the experiment, with substantial initial encystment that declined to near-zero by the third week of the experiment, when most potential infections had either been established or prevented (Figure 2A; plotted by population in Figure S3). Over the first two weeks, encystment trajectories differed by geographic region and were not affected by the early heatwave treatment (Table 1, Tables S4-S5). The northern group had high starting encystment levels (mean = 0.89, SD = 0.18) that declined steeply (−56%) by the next checkpoint. In contrast, initial encystment in the southern group was was moderate (mean = 0.65, SD = 0.31) and remained constant over the first two weeks. As described below, reduced and slower encystment was followed by limited infection in both the main southern population group and in the two excluded populations (supplemental results in Tables S6-S7), suggesting that less infectious parasite populations may generally be slower to recognize, attack, and/or penetrate into hosts.

**Figure 2.**
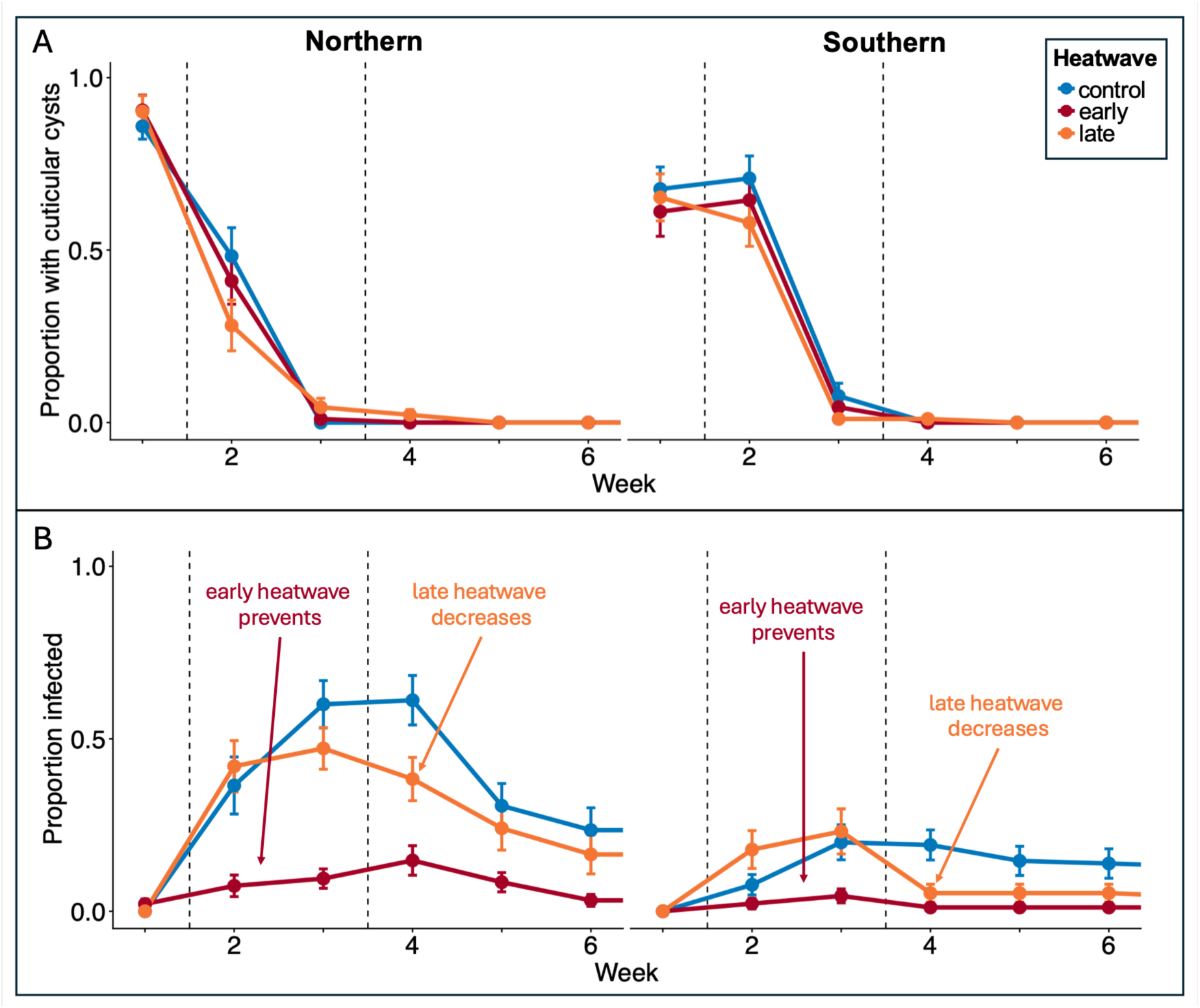
Parasite attack (encystment) varied regionally but was not affected by heat exposure. In contrast, heatwaves reduced infection regardless of geographic region and timing. In each panel, data from the first six weeks of the experiment are shown with the northern region at left and southern at right. Blue lines and points denote the temperature control groups, red denotes early heatwave groups, and orange denotes late heatwave groups. Points show mean values and error bars show +/-1 SE. Vertical dashed lines show when the early and late heatwave treatments were imposed. **(A)** Proportions of hosts with attacking parasites forming cuticular cysts; **(B)** Proportions of hosts with active or fatal internal parasite infections.

**Table 1.** Key statistically significant effects (p < 0.05) of heatwave treatment, geographic region, experiment checkpoint, and their interactions on the proportion of hosts attacked by, infected by, and mounting a melanization immune defense against parasites.

| GLMM key results |  |  |  |  |  | emmeans key results |  |  |  |  |
|---|---|---|---|---|---|---|---|---|---|---|
| Response | Predictors | Estimate | SE | Z-statistic | p-value | Contrasts | Estimate | SE | Z-ratio | p-value |
| Encystment | check 2 | -0.97 | 0.24 | -4.02 | <0.001 | northern vs. southern, check 1 | 1.52 | 0.47 | 3.23 | 0.001 |
|  | region | -0.62 | 0.25 | -2.53 | 0.01 | check 1 vs. 2, northern | 2.29 | 0.34 | 6.84 | <0.001 |
|  | check 2 x region | 1.08 | 0.24 | 4.5 | <0.001 |  |  |  |  |  |
| Infection | early heatwave | -2.48 | 0.41 | -6.02 | <0.001 | late heatwave vs. control, check 4 | -1.42 | 0.44 | 3.23 | 0.003 |
|  | region | -1.08 | 0.24 | -4.58 | <0.001 |  |  |  |  |  |
|  | late heatwave x check 4 | -1.0 | 0.27 | -3.63 | <0.001 |  |  |  |  |  |
| Immune melanization | late heatwave x check 4 | 0.92 | 0.25 | 3.65 | <0.001 | early heatwave vs. control, check 2, northern | 1.82 | 0.45 | 4.05 | <0.001 |
|  | early heatwave x region | -0.79 | 0.24 | -3.34 | <0.001 | late heatwave vs. control, check 4, northern | 1.37 | 0.49 | 2.81 | 0.01 |
|  | early heatwave x check 3 x region | 0.73 | 0.23 | 3.14 | 1.69 x 10<sup>-3</sup> |  |  |  |  |  |
|  | late heatwave x check 4 x region | -0.53 | 0.25 | -2.12 | 3.39 x 10<sup>-2</sup> |  |  |  |  |  |

Table note: Key results from generalized linear mixed effects models and emmeans post-hoc comparisons with Bonferroni-adjusted p-values. Encystment was modeled as a beta-binomial distribution with glmmTMB. Infection and melanization were modeled as binomial distributions with lme4. SE = Standard Error. Full results are presented in Tables S4-S5 (encystment), S8-S9 (infection), and S10-S11 (melanization).

### Heatwave effects on infection

Infection increased and then declined over time as larval hosts cleared infections or pupated, and parasite attack slowed (Figure 2B; plotted by population in Figure S4). Northern populations had greater infection rates than southern populations (northern: mean = 0.31, SD = 0.3; southern: mean = 0.11, SD = 0.19), parallelling previously observed field patterns. However, for both regions, heatwaves reduced infections compared to the control group at the next checkpoint (Table 1, Tables S8-S9). The early heatwave reduced the proportion of infected mosquitoes (at check 2, temperature control: mean = 0.28, SD = 0.3; early heatwave: mean = 0.06, SD = 0.12). The late heatwave also reduced the proportion of infected mosquitoes (at check 4, temperature control: mean = 0.36, SD = 0.33; late heatwave: mean = 0.21, SD = 0.26), but to a lesser extent that was somewhat sensitive to model specification (see *Robustness checks* below). Overall, heatwaves reduced infection regardless of population histories of exposure to winter heatwaves in their source environments, but the earlier heatwave had a stronger, more consistent effect.

### Heatwave effects on host defenses

#### 1. Melanization immune response

Paralleling the infection dynamics, the proportion of hosts mounting a melanization immune response to parasite attack initially increased before stabilizing and then declining (Figure 3A; plotted by population in Figure S5). Unlike heatwave effects on infection, impacts on melanization were only apparent for northern hosts (Table 1, Tables S10-S11). For the northern region, the early heatwave more strongly increased melanization than the late heatwave: compared to the temperature control, melanization was 117% higher for the early heatwave at the second checkpoint, and 84% higher for the late heatwave at the fourth checkpoint. Because we measured melanization as a binary outcome, and individual melanization spots may remain visible on larvae for multiple weeks, the observed change in melanization after the late heatwave represents a conservative estimate. In summary, heatwaves increased the melanization immune response only for the northern hosts that also experienced higher, quicker parasite attack and more infection overall.

**Figure 3.**
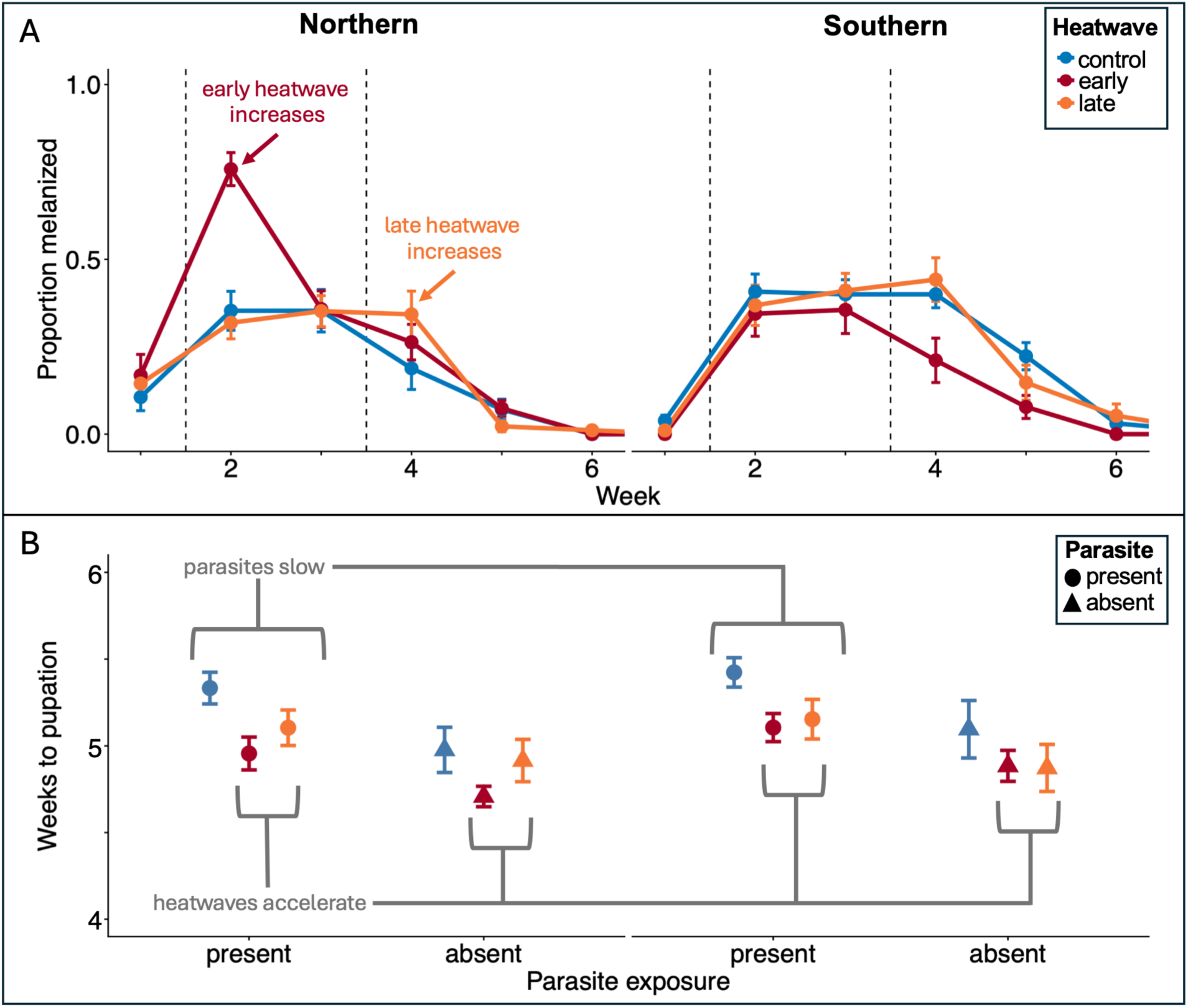
Heatwaves protected Northern hosts from parasites by increasing immune melanization, and hosts from both regions by accelerating development. In both panels, temperature control groups are shown in blue, early heatwave groups in red, and late heatwave groups in orange; northern region data are on the left and southern region data on the left; points show mean values and error bars show +/- 1 SE. **(A)** The proportions of parasite-exposed hosts exhibiting a melanization immune response in each temperature treatment group and geographic region, over the first six weeks of the experiment. **(B)** Host time to pupation by heatwave treatment, parasite exposure treatment, and region. Circles denote parasite exposure (infection treatment) and triangles denote parasite absence (control treatment).

#### 2. Larval development rate

Parasite exposure and heatwave treatments had opposing effects on the host’s larval development rate. For all hosts exposed to parasites, pupation occurred later (parasite present: mean = 5.19 weeks to pupation, SD = 0.7; parasite absent: mean = 4.92, SD = 0.73) (Figure 3B; plotted by population in Figure S6) (Table 2, Tables S12-13). Heatwaves counteracted the slowing effect of parasites and reliably sped up development (temperature control: mean = 5.27 weeks to pupation, SD = 0.75; early heatwave: mean = 4.92, SD = 0.6; late heatwave: mean = 5.01, SD = 0.75). These effects manifested as higher pupation hazard early on for heatwave-exposed hosts, and later for parasite-exposed hosts. Time to pupation did not differ significantly between the northern and southern regions. Overall, developmental acceleration in response to heatwaves was general across heatwave timing, geographic regions, and parasite presence, with no statistically significant interactions between these predictors.

**Table 2.** Key statistically significant effects (p < 0.05) of experiment checkpoint, parasite exposure, and heatwave treatment, and their interactions on host time to pupation and survival to pupation.

|  | GLMM key results |  |  |  |  | emmeans key results |  |  |  |  |
|---|---|---|---|---|---|---|---|---|---|---|
| Response | Predictors | Estimate | SE | Z-statistic | p-value | Contrasts | Estimate | SE | Z-ratio | p-value |
| Host development rate | Check 5 | 2.75 | 0.24 | 11.25 | <0.001 | Early heatwave - control, check 4 | 0.85 | 0.24 | 3.56 | <0.001 |
|  | Check 6 | 3.93 | 0.28 | 13.90 | <0.001 | Late heatwave - control, check 4 | 0.8 | 0.24 | 3.29 | 2 x 10<sup>-3</sup> |
|  | Check 7+ | 4.02 | 0.50 | 8.09 | <0.001 | Early heatwave - control, check 5 | 0.65 | 0.16 | 4.1 | <0.001 |
|  | Parasite absent | 1.41 | 0.27 | 5.33 | <0.001 | Late hatwave - control, checks 7+ | -2.13 | 0.76 | -2.81 | 0.01 |
|  | Early heatwave | 1.09 | 0.29 | 3.76 | <0.001 | Parasite present - absent, check 4 | -1.15 | 0.19 | -6.06 | <0.001 |
|  | Late heatwave | 0.95 | 0.30 | 3.23 | 1.22 x 10<sup>-3</sup> | Parasite present - absent, check 5 | -0.29 | 0.13 | -2.24 | 0.03 |
|  | Check 6 x<br>late heatwave | -1.22 | 0.33 | -3.65 | <0.001 | Parasite present - absent, check 6 | -0.71 | 0.24 | -2.94 | 3.3 x 10<sup>-3</sup> |
|  | Check 7+ x<br>late heatwave | -2.02 | 0.63 | -3.23 | 1.26 x 10<sup>-3</sup> |  |  |  |  |  |
|  | Check 5 x<br>parasite absent | -0.85 | 0.20 | -4.21 | <0.001 |  |  |  |  |  |
|  | Check 7+ x<br>parasite absent | -1.91 | 0.75 | -2.53 | 0.01 |  |  |  |  |  |
| Host survival | Early heatwave | 1.35 | 0.29 | 4.61 | <0.001 | early heatwave - control, parasite present | -1.35 | 0.29 | -4.61 | <0.001 |
|  | Parasite absent | 2.44 | 0.49 | 4.98 | <0.001 | parasite present - absent, late heatwave | -2.14 | 0.49 | -4.32 | <0.001 |
|  | Early heatwave<br>x parasite<br>absent | -1.50 | 0.69 | -2.18 | 0.03 | parasite present - absent, no heatwave | -2.44 | 0.49 | -4.98 | <0.001 |

Table note: GLMM key results from generalized linear mixed effects models; emmeans key results from post-hoc pairwise comparisons with Bonferroni-adjusted p-values; SE = Standard Error. Full results are presented in Tables S12-S13 (development rate) and S14-S15 (survival).

### Heatwave and parasite effects on host survival

Although both heatwaves accelerated host development and decreased infection, only the early heatwave rescued hosts from parasite-induced mortality, recovering pupation to the high levels observed when parasites were absent (across all temperature treatments, without parasites: mean proportion surviving = 0.96, SD = 0.09; with parasites and early heatwave, mean = 0.9, SD = 0.14) (Figure 4; plotted by population in Figure S7) (Table 2, Tables S14-15). Survival to pupation was lower for mosquitoes exposed to parasites and either no heatwave (mean proportion surviving = 0.71, SD = 0.27) or the late heatwave (mean proportion surviving = 0.73, SD = 0.23). Heatwaves in the absence of parasite exposure did not affect host survival, and survival outcomes did not vary by geographic region. In summary, only early heatwaves before infections had taken hold conferred a survival advantage to hosts exposed to parasites, which otherwise decreased host survival to pupation.

**Figure 4.**
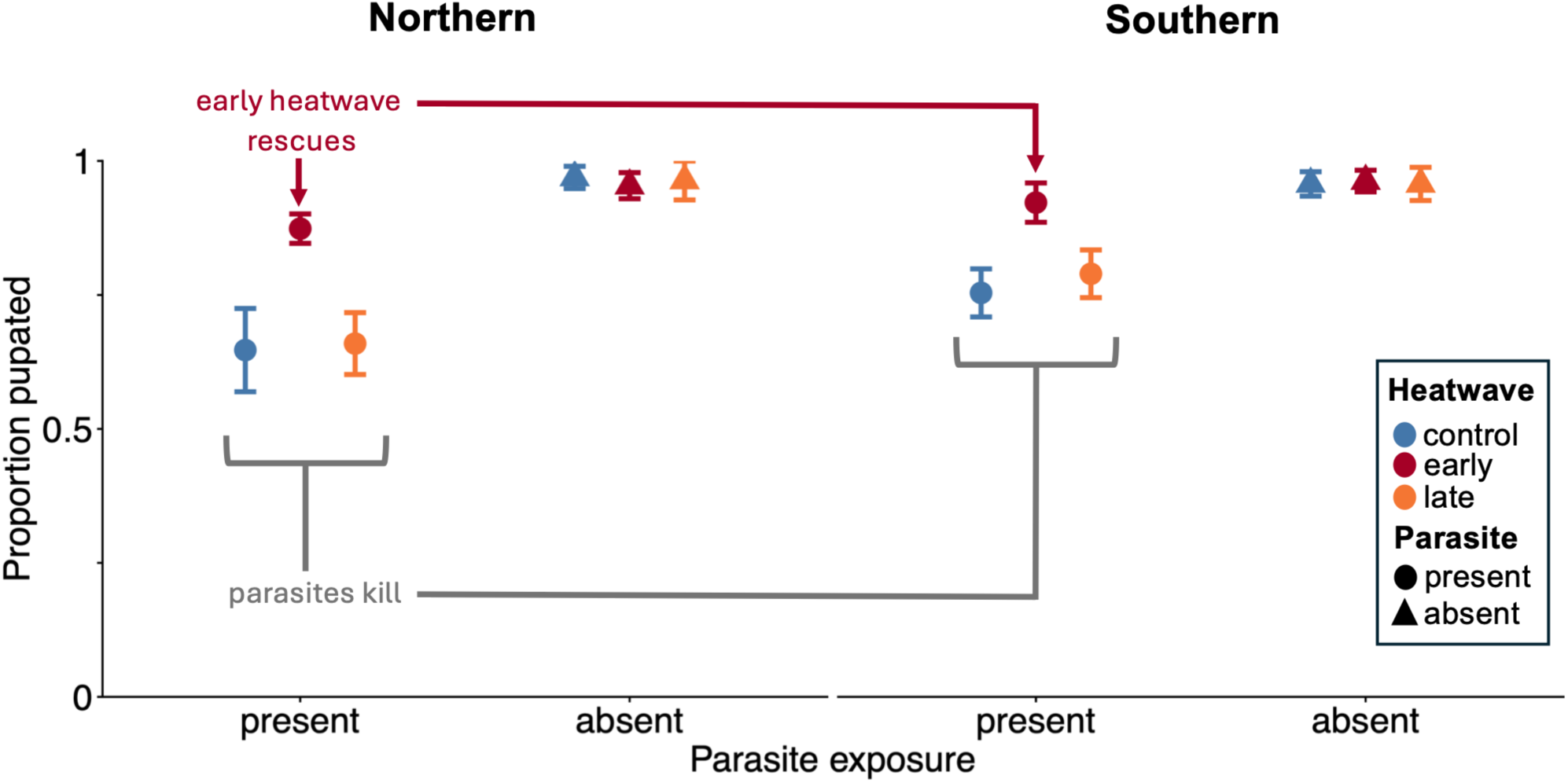
Parasite exposure reduced host survival, but the early heatwave had a rescuing effect. The final proportions of hosts surviving to pupation are shown by temperature treatment, parasite exposure, and geographic region. Colors correspond to heatwave treatments (blue = control, red = early heatwave, orange = late heatwave); parasite presence is indicated by circles and absence by triangles; points show group means, with error bars spanning +/- 1 SE. Outcomes are shown for the northern region on the left, and the southern region on the right.

### Robustness checks

The robustness of the results to variation among populations within geographic regions was broadly supported by supplemental models that replaced region fixed effects with population ID (encystment: Tables S16-S19; infection: Tables S20-S23; melanization: Tables S24-S27); development speed: Tables S28-S31; survival: Tables S32-35). However, there were some exceptions. First, the late heatwave reduction in infections was not detected in a robustness check that modeled three-way interactions between heatwave treatment, checkpoint, and population ID (Table S16). Higher model complexity, a relatively weak reduction in infections, and limited replication for testing the three-way interaction likely contributed to this difference from the regional analysis and a simpler robustness check that included population ID only as an additive effect (Table S18). Next, although time to pupation did not vary geographically when models interacted region (main analysis) or population ID with heatwave treatment and checkpoint (Table S28), significant variation among populations within regions was detected when population was included only as a main effect (Table S30). However, heatwave and parasite exposure effects on larval development speed were consistent across all three analyses. Finally, both robustness checks for host survival identified the Alameda parasite population as associated with particularly high host mortality, but heatwave and parasite exposure effects were again consistent for all three models (Figure S7, Tables S32, S34).

## Discussion

In this experiment, the effects of short heatwaves on parasitism at different times in the ontogeny of infection were mechanistically predictable from combined knowledge of host – parasite thermal biology and natural history. In a winter-adapted study system, heatwaves reduced infection (Figure 2B) and increased host defenses (Figure 3), consistent with the cool thermal optima for infection and elevated immune and developmental defenses at warm constant temperatures identified in previous TPC experiments (Figure 1B) (Couper et al. 2024; Farner et al. 2026). Early heatwaves were particularly beneficial for hosts (Figure 4) because they increased host defenses during the parasite attack stage and prevented infection. However, counter to our hypothesis of local adaptation to heatwaves, populations with different histories of winter heat exposure (Figure 1C) did not differ in their infection responses to heatwaves, which were consistent despite regional variation in climate, parasite infectiousness, and host immune melanization sensitivity. Overall, the results demonstrate the importance of incorporating heat anomalies and their timing into assessments of temperature-dependent disease risk.

The results add to evidence that heatwaves can drive parasitism with effects that are complex, yet predictable based on fundamental knowledge of temperature-dependent host and parasite traits. The reduced infection we observed following 24°C heatwaves aligned with our previous observations of low infection probability at warm constant temperatures (Farner et al. 2026; Lyberger et al. 2024). Elevated host defenses following heatwaves were also consistent with warm thermal optima for host immune melanization and development rates in previous TPC experiments (Couper et al. 2024; Farner et al. 2026). While parasite exposure slowed host development, heatwaves rescued host development rates. In contrast, although warm constant temperatures were previously associated with lower parasite encystment, the first stage of attack (Lyberger et al. 2024), short-term heat exposure did not observably affect this trait. This may be explained by heat exposure occurring after cuticular cyst formation; slower thermal acclimation of this parasite trait; or a stronger influence of genetics driving *L. clarki* attack.

Importantly, heatwave timing within the host – parasite interaction determined impact. Although both heatwaves reduced infection compared to the temperature control, only the early heatwave improved host survival. When host defenses were strengthened by early heatwaves during parasite attack, most infections were prevented, and host survival increased as expected from TPCs. While the late heatwave accelerated host development commensurately and was also followed by increased melanization in the northern hosts, it caused a smaller decrease in infections and did not change host survival. Thus, brief heat exposure could not fully shift the balance towards hosts once parasite infections were established, intensifying, and depleting host resources. We did not track heatwave impacts beyond pupation, but downstream effects on host and parasite population dynamics might also be consequential. For example, heatwaves and their timing might affect adult mosquito longevity, body size, and body quality, and by extension, the reproductive capacity, offspring quality, and dispersal of both hosts and parasites surviving to later stages (Pottier *et al*. 2026). More broadly, it is important to test whether combined knowledge of TPCs and natural history can clarify the seemingly idiosyncratic heatwave effects documented across the disease ecology literature. For most systems, we lack knowledge of the thermal biology of key host and parasite traits that govern the infection process, and characterizing relevant TPCs will be a crucial first step.

Host and parasite local thermal adaptation may determine the geography of current and future temperature-dependent disease risk, but whether historical climate conditions shape heatwave responses is rarely assessed. We found that heatwave impacts on parasitism and host survival did not vary based on historical winter heat exposure in northern versus southern populations. For both regions, parasite exposure slowed development, and heat exposure counteracted this effect by shortening time windows for encysted parasites to enter larvae (early heatwave), and for internal infections to replicate to lethal densities (both heatwaves). These results suggest similar responses across the large species range, a pattern we have also observed in previous experiments comparing constant temperature effects among *Ae. sierrensis* and *L. clarki* populations (Couper et al. 2024; Farner et al. 2026; Lyberger et al. 2024). The few other studies investigating intraspecific variation in parasitism responses to heatwaves or constant temperatures also found no evidence of local adaptation (Landis et al. 2012, Franke et al. 2019).

Interestingly, the uniform effects of heatwaves on infection and host survival arose despite geographic variation in some host and parasite traits. Heatwaves increased the melanization immune response only for northern region hosts with low historical winter heat exposure. Whether this result corresponds to host local adaptation to parasites, heatwaves, or neither is unclear because these hosts also experienced higher levels of parasitism, whereas the southern group’s lower parasite attack may have masked a similar melanization response. Geographic differences in parasite traits also did not modify heatwave impacts on infection. Although previous work showed that free-living *L. clarki* ciliate thermal optima are locally adapted (Ismail et al. 2023; Lyberger et al. 2024), our finding of consistent heatwave impacts across regions instead matches the uniform constant and mean temperature effects on parasitism previously observed over 11° latitude of the species range (Farner et al. 2026).

Heatwave effects on infection were also maintained across substantial variation in parasite infectiousness on sympatric hosts. However, a near absence of infection precluded heatwave impacts for the Marin and Los Angeles county populations. In a supplementary experiment investigating the low infectiousness of these two parasite populations, both were able to infect allopatric hosts, implying that host local adaptation to these parasites may have prevented infection in the main heatwave experiment (Supplementary text and Figures S9-S10). In contrast, we have previously observed widespread parasite local adaptation to sympatric hosts, including for the Marin county population (and two others) used in both studies (Lyberger et al. 2024). Collectively, these results suggest that geographic and interannual variation in parasite attack and host susceptibility might affect the overall infection rate, while heatwave impacts on parasitism remain consistent, highlighting the interplay between climate gradients, interannual weather variation, and host – parasite coevolution as a future research direction.

Although the results of our lab experiment aligned with our previous knowledge of temperature effects on this host – parasite interaction from multiple experiments and a multiyear, large-scale field survey in this study system (Ismail et al. 2023; Farner et al. 2026; Lyberger et al. 2024), further research is needed to assess links to outcomes in the field. Natural temperature patterns are comparatively variable, and many characteristics of experimental temperature treatments other than heatwave timing are known to affect parasitism (Kunze et al. 2022; McCartan et al. 2025; Studer & Poulin 2013; Taig et al. 2025; Rozmann et al. 2026). Additionally, abiotic and biotic factors such as tree hole habitat buffering against extreme air temperature fluctuations; temperature and humidity levels during host egg and parasite resting stages; and opportunities for behavioral thermoregulation might shape outcomes under extreme heat (Boutry et al. 2026; Marcus et al. 2023; Porras et al. 2023). Understanding of how heatwave effects are shaped by these and other factors will require long-term field surveillance and experiments that relate natural and artificial heatwaves to parasitism in more complex habitats.

### Conclusions

Ongoing warming is widely expected to reshape disease risk in nature (Altizer et al. 2013; Rohr & Cohen 2020). For the *Ae. sierrensis* – *L. clarki* system, our results suggest that increased exposure to winter heatwaves that reduce parasite fitness may accelerate distributional shifts in parasitism compared to gradual warming alone. The southern range edge parasite populations in this study appeared particularly vulnerable, as they were poorly adapted to cope with both their hosts and with warm temperatures that regularly occur in their source climates. The extent to which winter heatwaves drive geographic shifts in parasite pressure will also depend on how interannual variation in heatwave occurrence shapes host – parasite coevolutionary dynamics, particularly the maintenance of host resistance (Li *et al*. 2025). Assessments of current and future disease risk therefore need to consider not just changes in average temperatures, but also in the frequency, intensity, and timing of extreme temperatures (Claar & Wood 2020). Our findings highlight that fundamental knowledge of host – parasite thermal biology and natural history can provide insight into this complex problem.

