## Supplemental Information for "Heatwaves rescue a mosquito host from parasitism across a large geographic gradient"

### Methods and Results

1. *Weather station data analysis*

To design treatments simulating ecologically relevant winter temperature conditions and heatwaves, we analyzed 35 years of PRISM daily temperature and precipitation data (1988 – 2023) for the field locations from which the experimental populations were sourced (PRISM Climate Group and Oregon State University 2021)

First, to determine the relevant time period for our temperature analyses, we used precipitation data to determine when during the winter rainy season in California tree hole mosquito and ciliate activity has been most likely to begin over the long term. *Aedes sierrensis* larvae and free-living *Lambornella clarki* ciliates have previously been observed to appear within 24 hours of tree hole habitat filling with water (Washburn et al. 1991). In our field observations spanning November 2021 – March 2025, mosquito hatching has first occurred in late December or early January 75% of the time, and required approximately 50 – 80 mm rainfall over a short time span of 1-2 weeks. Based on these observations, we used PRISM data to estimate the long-term seasonality of tree hole habitats in the study area by calculating mean precipitation at quarter-monthly intervals for November, December, and January at each field site over the past 35 years. These data showed that, on average, the period between mid-December and mid-January has rainfall sufficient for tree hole filling in both southern and northern California, in alignment with our recent field observations (although northern tree holes often fill about one week earlier) (Figure S8). Based on these observations, we used long-term January temperature data to design experimental temperature regimes.

To select a baseline daily temperature fluctuation regime, we calculated the mean daily maximum and minimum January temperatures across all population source sites. These were 16ºC and 6ºC, respectively. We set the baseline daytime incubation temperature of 17ºC and nighttime temperature of 7ºC (the coldest achievable temperature with our incubators) (Figure S9).

To select ecologically relevant heatwave treatment temperatures, we analyzed the long-term daily maximum and minimum temperature data for the population source locations. To identify a temperature that would be both realistic and unusually hot for all populations, we first estimated the 97.5$^{th}$ percentile January daily maximum temperature for each field site as the mean daily maximum temperature plus 2 SDs. This ranged from 19.5ºC for Marin County to 28.4ºC for Los Angeles County. We calculated the mean of these estimated natural heatwave temperatures as 23.8ºC and then used the PRISM time series data for each field site to confirm that this temperature had occurred at least once from 1988 to 2023. Because the hottest January temperatures recorded for each site ranged from 24ºC (Alameda) to 32.3ºC (Los Angeles), we set the daytime heatwave incubation temperature to 24ºC. To choose the heatwave nighttime incubation temperature, we calculated the mean daily minimum temperature for January days with daily maximum temperatures between 22ºC and 26ºC at each site, and across all sites. The mean daily low temperatures associated with hot January days ranged from 5.4ºC (Marin) to 12.6ºC (Los Angeles), with an overall mean of 7.4ºC. Therefore, we maintained the baseline 7ºC nighttime incubation temperature during our heatwave treatments.

Finally, to contextualize the heatwave treatment temperatures, we plotted the number of January days at each field site exceeding 23ºC over time (Figure 2), and estimated the percentile into which 24ºC fell among all daily maximum temperatures for each site. Summaries of the PRISM temperature analyses are provided in Table S2. PRISM data were extracted using Google Earth Engine and analyzed in R version 4.2.1 (Gorelick et al. 2017; R Core Team 2022).

2. *Statistical analyses: Checking robustness of regional model results at the population level*

In order to assess the extent to which early and late heatwave effects varied among populations within climatic regions, we fit two population-level models as robustness checks. The first model tested for complex three-way interactions between heatwave treatment, checkpoint, and population, and included well ID as a random effect. Because the overall amount of infection differed substantially among populations and replication was low for some population x heatwave treatment groupings, this model was limited in its power to detect interactive effects of heatwave treatment and checkpoint. A second robustness check focused more directly on the interaction between heatwave treatment and checkpoint by simplifying population ID to an additive effect.

For some models that included complex three-way interaction terms, complete separation occurred and we used the R package "blme" (Dorie & Dorie 2015) to set weak priors on fixed effects estimates, following the recommendations from Gelman et al. (2008). Because the population-level models for survival to pupation used only the final observation for each well and did not include any random effects, the three-way interaction model was fit using the function "bayesglm" from the R package "arm" (Gelman *et al.* 2015), and the two-way interaction model was fit using the base R function "glm".

3. *Experiment investigating low infectiousness of sympatric parasites*

*3.1 Experiment*

To investigate whether the very low infection rates in the Marin and Los Angeles County populations were due to weakly infectious parasite cultures or host resistance to sympatric parasites, we ran a small follow-up experiment with allopatric host – parasite pairs including these source populations. Using the same methods as the main experiment, we monitored infection rates under the baseline daily temperature fluctuation for three weeks for each of the following groups: Los Angeles county parasites paired with San Mateo (N = 4 wells), San Diego (N = 4 wells), and Los Angeles (N = 8 wells) hosts; Marin County parasites paired with San Mateo (N = 4 wells), San Diego (N = 4 wells), and Los Angeles (N = 8 wells) hosts; and Los

Angeles hosts paired with San Mateo (N = 8 wells), Santa Barbara (N = 8 wells), and Los Angeles (N = 8 wells) parasites. Each well contained 5 larvae. These pairings aimed to cover both geographic regions and the variation in infection rates observed in the main experiment to the extent possible with our remaining materials (e.g., there were sufficient remaining mosquito eggs for Los Angeles, but not for Marin County).

*3.2 Analysis*

To test for host local adaptation to parasites, we used binomial generalized linear mixed models in the R package glmmTMB (Brooks et al. 2017), with final infection rate as the dependent variable. For hosts exposed to the two low-infection parasite cultures, the model predictors included geographic distance between source locations (as a proxy for genetic relatedness to the sympatric population), parasite population ID, and their interaction. Host population was included as a random effect, and geographic distance values were scaled and centered. For pairings of the Los Angeles County host population with sympatric and allopatric parasites, we similarly modeled the effects of geographic distance of the parasite population on the final infection rate, with parasite population ID included as a random effect.

*3.3 Results*

Infection rates were positively correlated with geographic distance between parasite and host source sites, both when the two cultures with low sympatric infection rates were paired with allopatric hosts (estimated effect = 0.004, Z-Value = 1.98, p-value = 0.047) (Figure S9), and when the host population with low sympatric infection was paired with allopatric parasites (estimated effect = 0.006, Z-Value = 2.37, p-value = 0.017) (Figure S10). Paralleling the geographic pattern in parasite infectiousness in the main experiment, the southern region low-infection parasite culture from Los Angeles county had significantly lower infection rates overall compared to the northern region, low-infection parasite culture from Marin county (mean proportion infected by Los Angeles parasite = 0.06, sd = 0.12; Marin parasite mean = 0.27, SD = 0.22; estimated effect = -2.21, Z-value = -2.4, p-value = 0.016). However, Los Angeles and Marin county parasite infection levels were similar for geographically distant hosts, suggesting that the southern parasites were not inherently less infectious.

Finally, although the data suggested a stronger correlation for the Los Angeles compared to the Marin parasite culture, this interaction was not statistically significant (p = 0.05). Factors contributing to the weaker relationship between distance and infection for the Marin county parasite might include: the absence of this particular sympatric host – parasite pairing in the supplemental experiment due to insufficient remaining Marin mosquito eggs; differences in host resistance mechanisms underlying low sympatric infectiousness, and their geographic variation; and/or unknown differences in other experimental conditions between the main and supplemental experiment that affected the overall infectiousness of the Marin parasites.

**Figures**

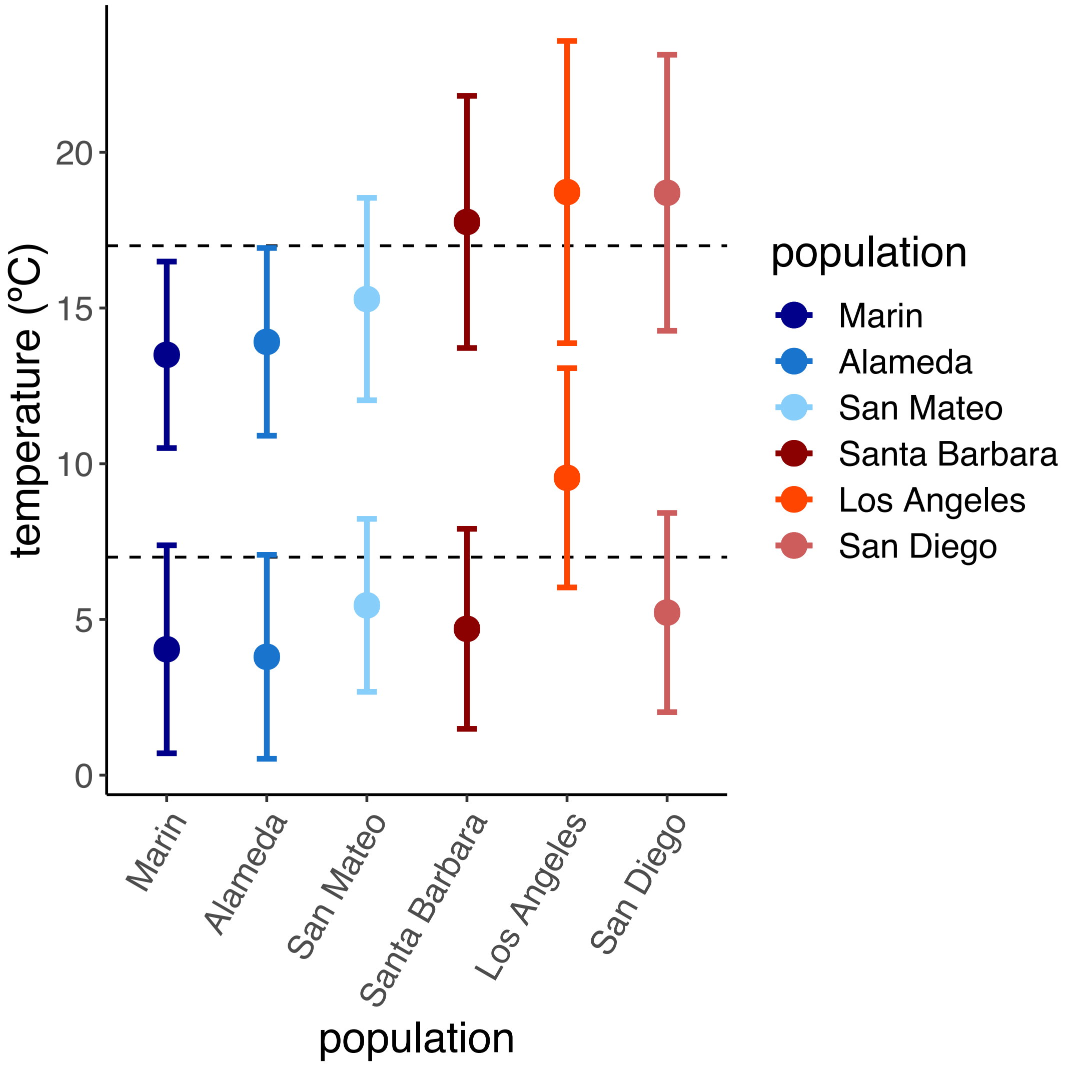


**Figure S1. Experimental baseline daytime and nighttime incubation temperatures (dashed lines) were intermediate to long-term mean daily maximum (top row) and minimum (bottom row) January temperatures across the locations from which the six experimental populations were sourced (colored points with error bars showing +/- 1 SD).**

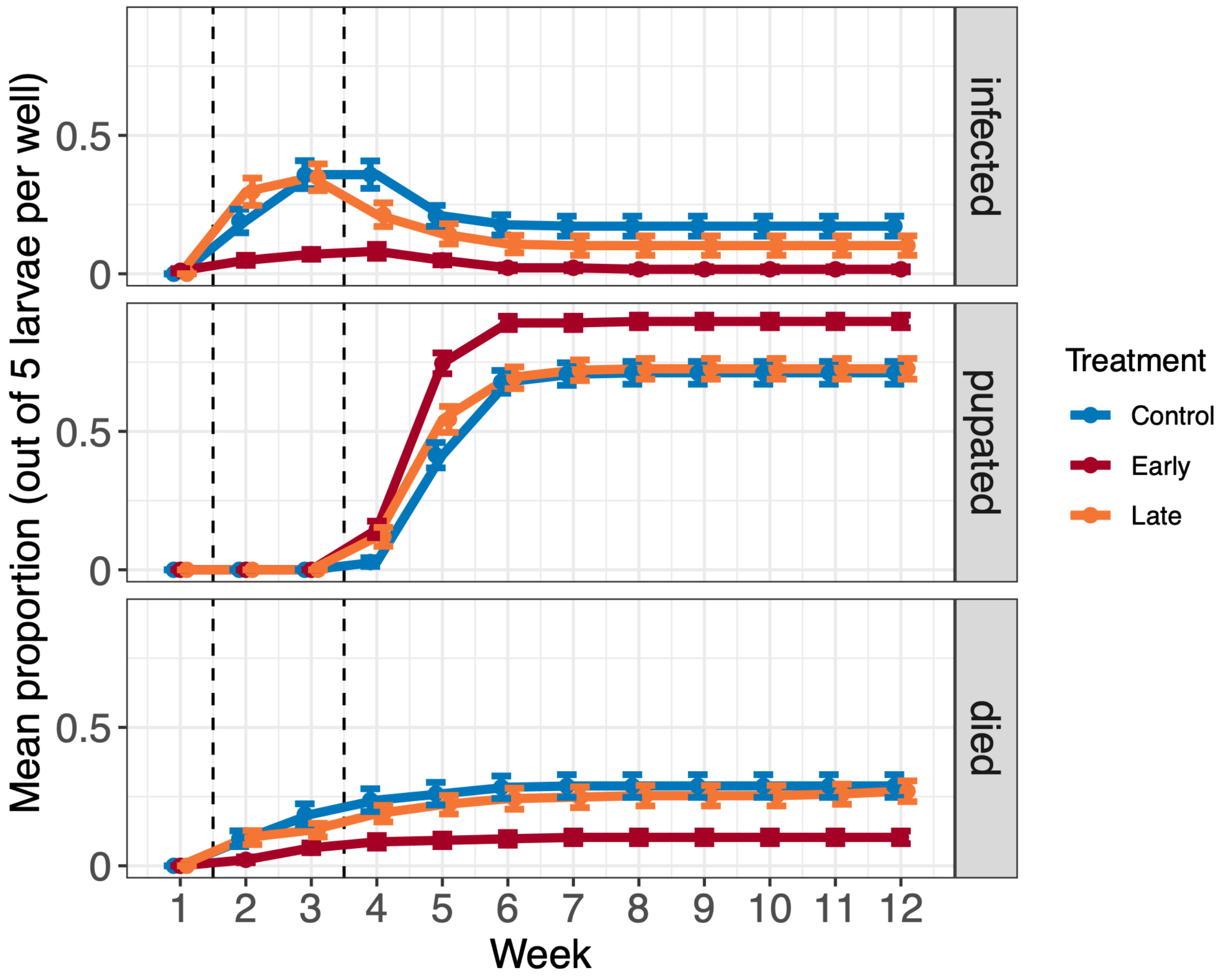


**Figure S2. Trajectories of infection, pupation, and host deaths throughout the experiment, by heatwave treatment, averaged across all populations.** Points show mean proportions for each outcome at each weekly checkpoint, error bars show +/- 1 SE, colors indicate heatwave treatments (blue = no heatwave, red = early heatwave, orange = late heatwave), and dashed lines show the timing of the two heatwave treatments.

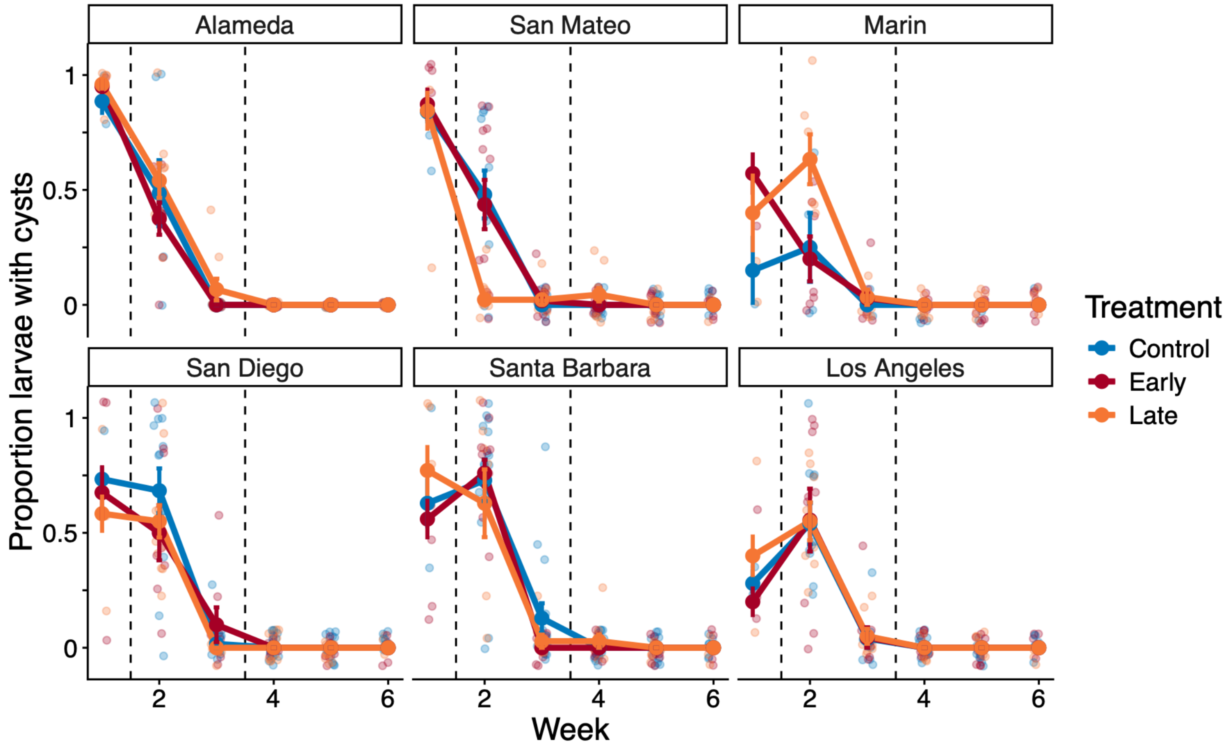


**Figure S3. Parasite attack rates over the first six weeks of the experiment, for each population and heatwave treatment.** The top row shows the northern populations, and the bottom row shows the southern populations. Colors correspond to heatwave treatments, with blue indicating no heatwave, red the early heatwave, and orange the late heatwave. Vertical dashed lines show when each heatwave treatment occurred. Solid points and lines track mean values, error bars show +/- 1 SE, and translucent points show data from individual microcosms housing five mosquito larvae each. The Marin and Los Angeles populations were excluded from all main analyses due to very low infection.

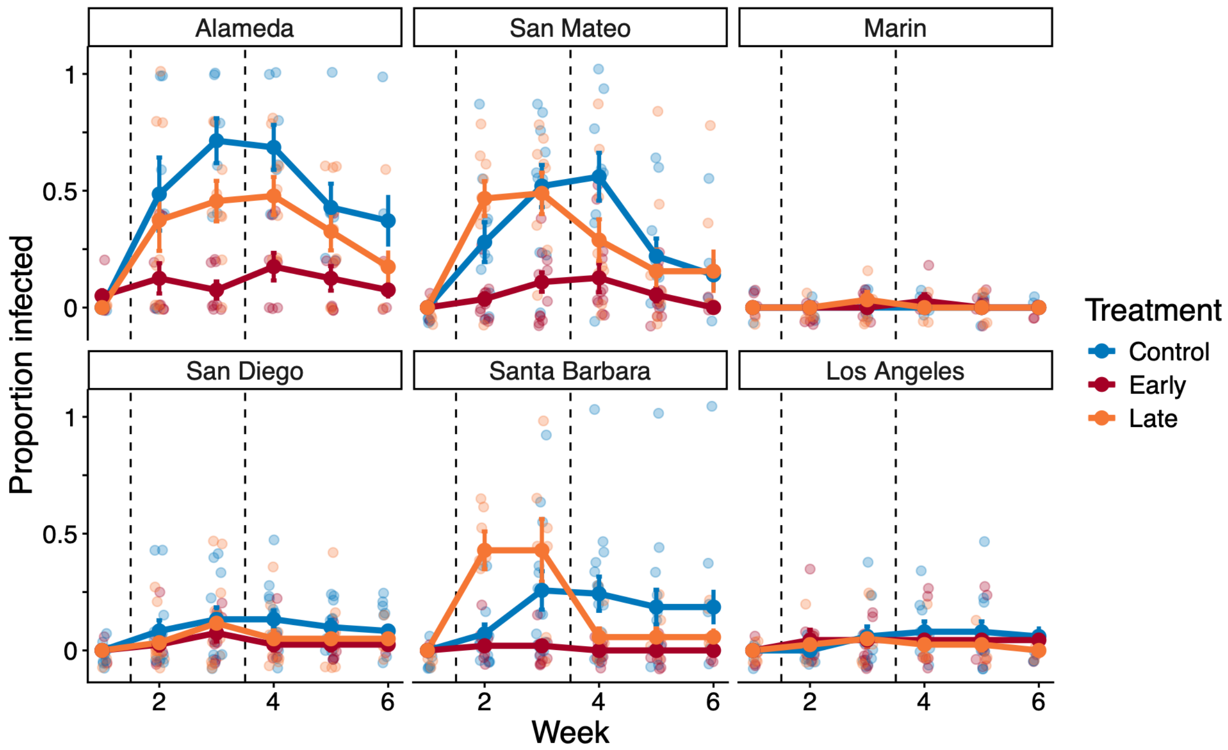


**Figure S4. Infection rates over the first six weeks of the experiment, for each population and heatwave treatment.** The top row shows the northern populations, and the bottom row shows the southern populations. Colors correspond to heatwave treatments, with blue indicating no heatwave, red the early heatwave, and orange the late heatwave. Vertical dashed lines show when each heatwave treatment occurred. Solid points and lines track mean values, error bars show +/- 1 SE, and translucent points show data from individual microcosms housing five mosquito larvae each. The Marin and Los Angeles populations were excluded from all main analyses due to very low infection.

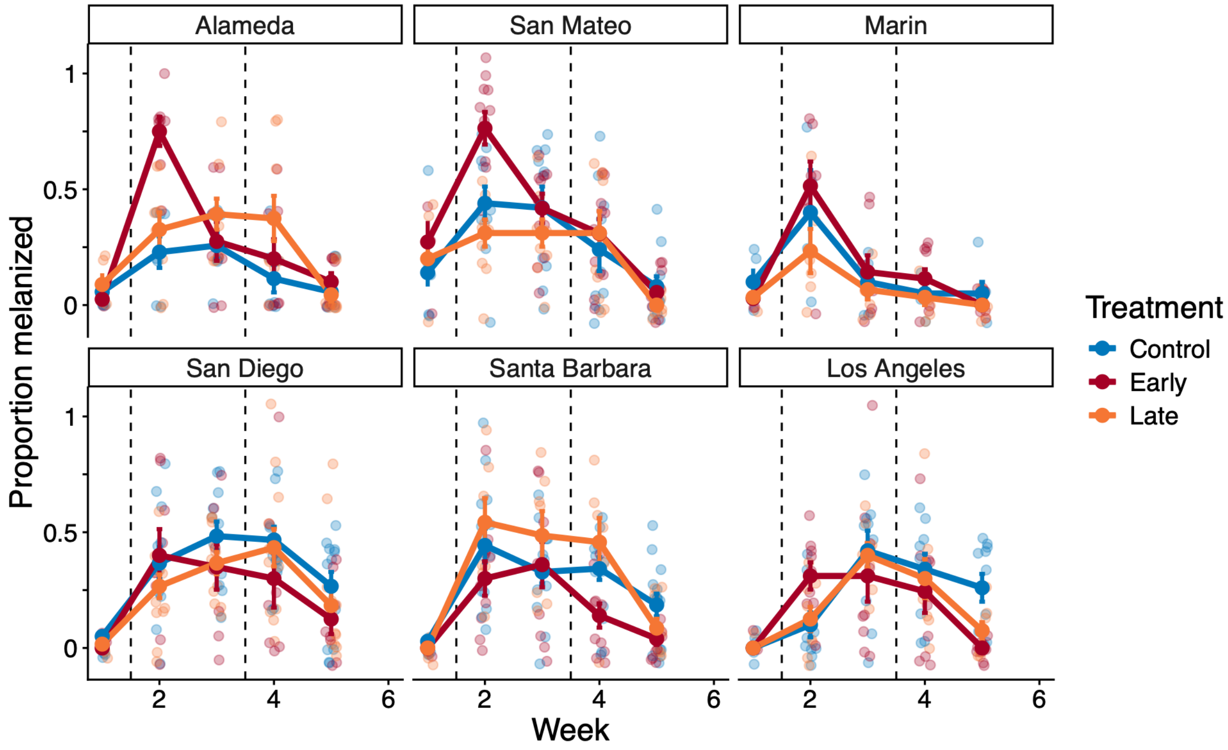


**Figure S5. Host melanization immune response rates over the first six weeks of the experiment, for each population and heatwave treatment.** The top row shows the northern populations, and the bottom row shows the southern populations. Colors correspond to heatwave treatments, with blue indicating no heatwave, red the early heatwave, and orange the late heatwave. Vertical dashed lines show when each heatwave treatment occurred. Solid points and lines track mean values, error bars show +/- 1 SE, and translucent points show data from individual microcosms housing five mosquito larvae each. The Marin and Los Angeles populations were excluded from all main analyses due to very low infection.

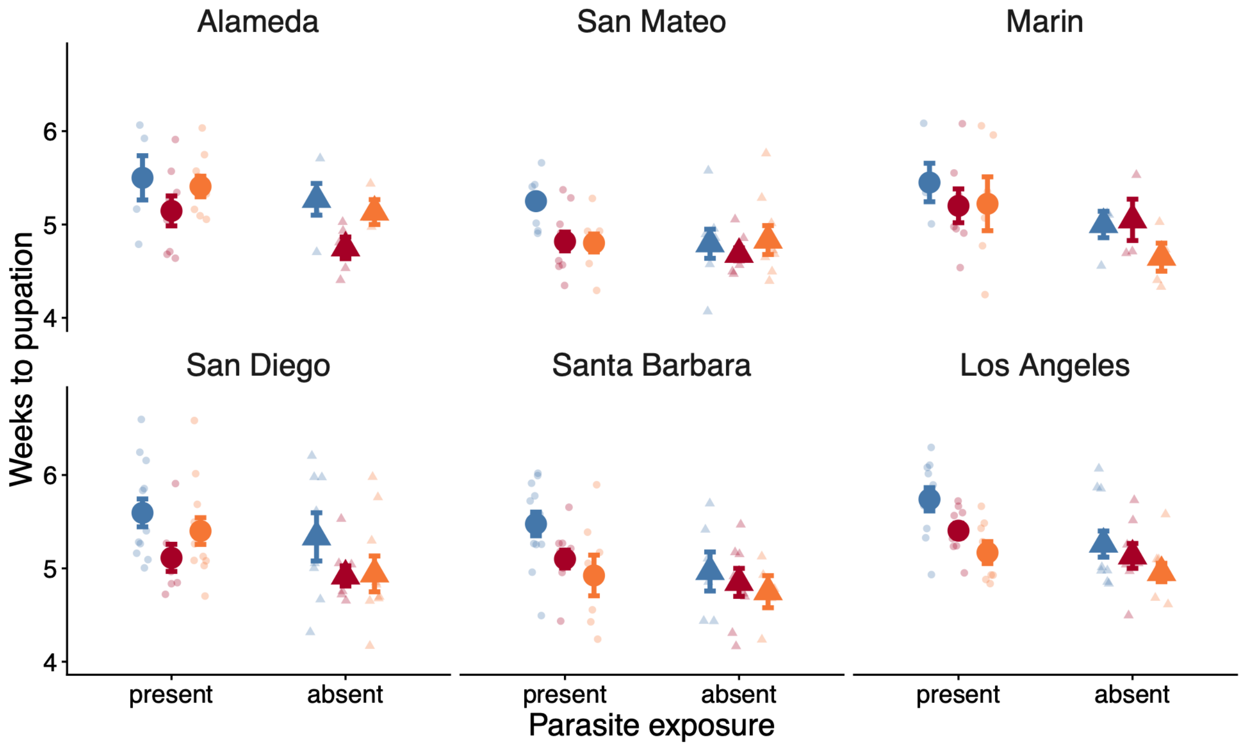


**Figure S6. Weeks to pupation for each population, by heatwave treatment and parasite exposure.** The top row shows the northern populations, and the bottom row shows the southern populations. Colors correspond to heatwave treatments, with blue indicating no heatwave, red the early heatwave, and orange the late heatwave. Solid points show mean values, error bars show +/- 1 SE, and translucent points show data from individual microcosms housing five mosquito larvae each. The Marin and Los Angeles populations were excluded from all main analyses due to very low infection.

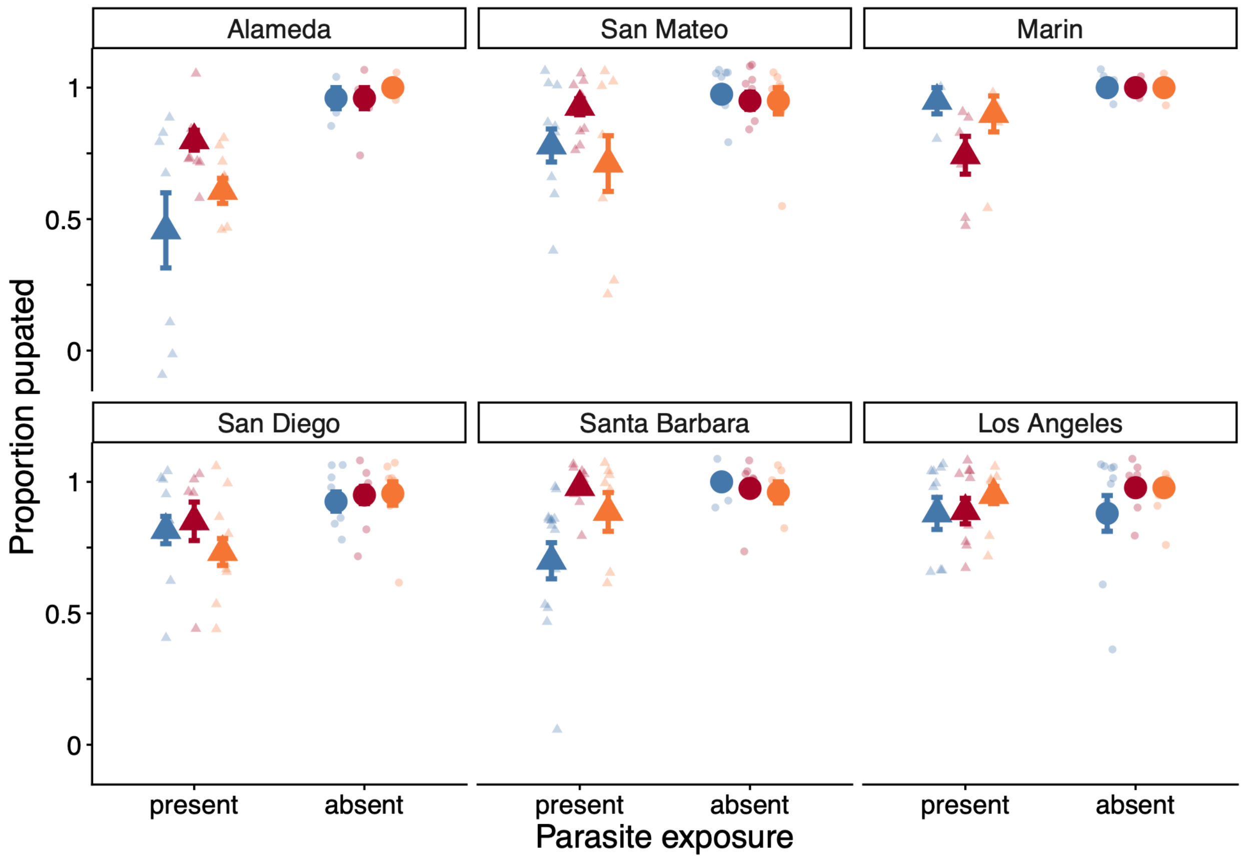


**Figure S7. Proportion mosquito larvae surviving to pupation for each population, by heatwave treatment and parasite exposure.** The top row shows the northern populations, and the bottom row shows the southern populations. Colors correspond to heatwave treatments, with blue indicating no heatwave, red the early heatwave, and orange the late heatwave. Solid points show mean values, error bars show +/- 1 SE, and translucent points show data from individual microcosms housing five mosquito larvae each. The Marin and Los Angeles populations were excluded from all main analyses due to very low infection.

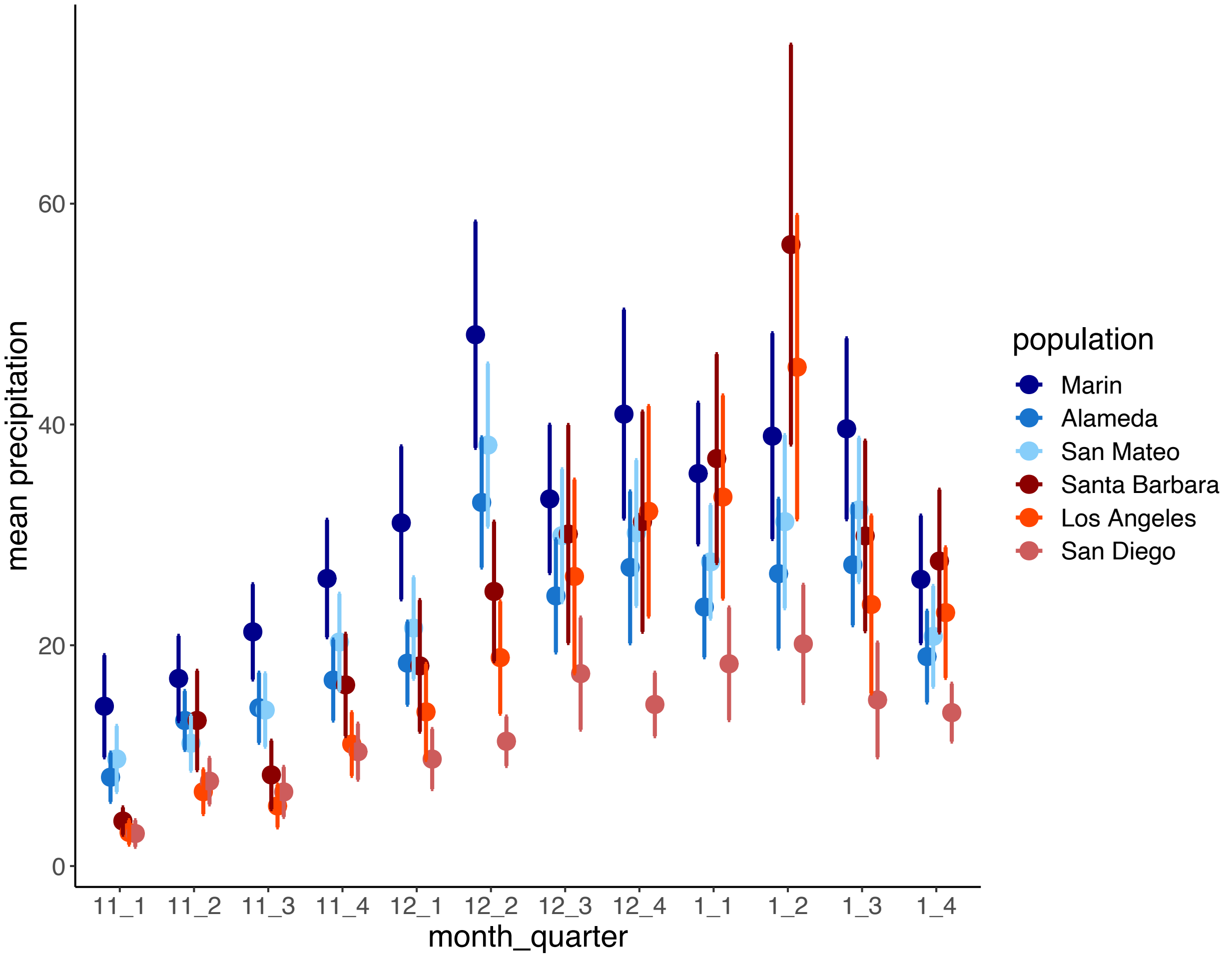


**Figure S8. Sufficient rainfall for tree hole habitat filling across the study area was most likely to occur between mid-December and mid-January between 1988 and 2023.** Each point shows the long-term mean quarter-monthly precipitation estimate for a field site from which an experimental population was sourced, calculated from PRISM data. Error bars show +/- 1 SE. Northern California sites are shown in shades of blue and southern sites in shades of red.

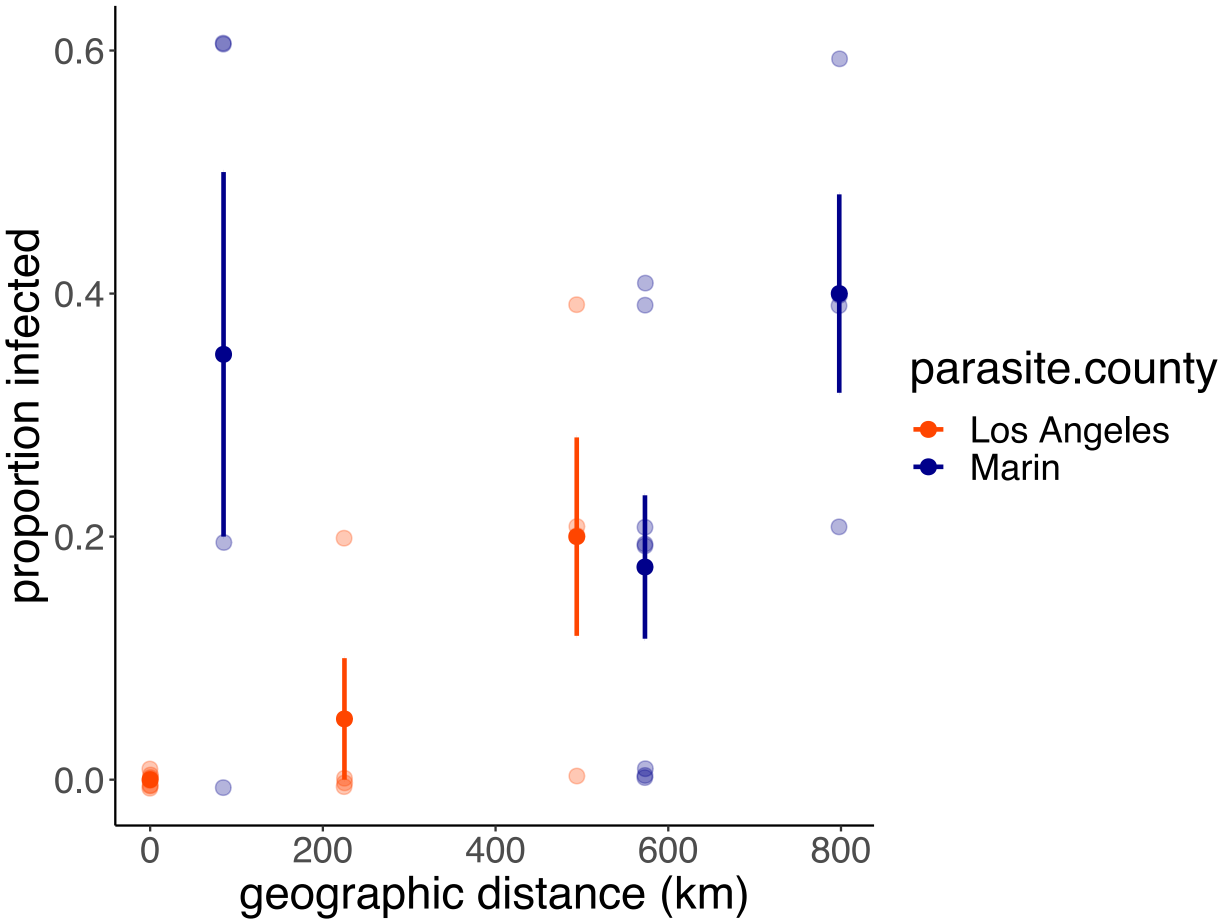


**Figure S9. The two parasite cultures with very low infection rates in the main experiment were more infectious on allopatric hosts from distant locations.** Solid points show mean values for a given host – parasite population pairing, with error bars showing +/- 1 SE. Data are colored by parasite population, with orange indicating the Los Angeles County parasite culture, and blue indicating Marin County. Translucent points show data for individual microcosms containing 5 mosquito larvae each. The Los Angeles parasite culture was less infectious overall than the Marin culture ($p = 0.02$).

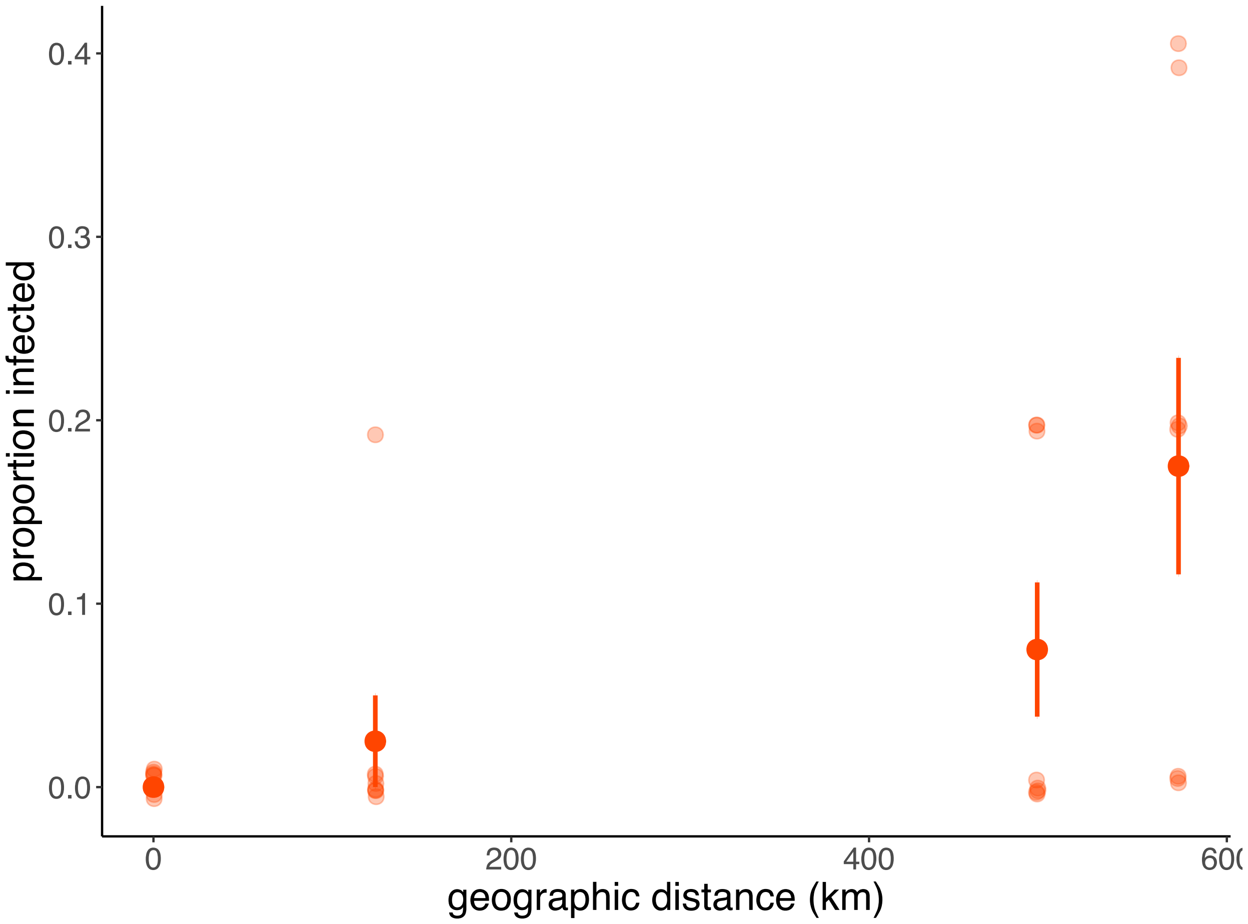


**Figure S10. The Los Angeles County host population with very low infection rates in the main experiment was more susceptible to infection by allopatric parasites from farther away.** Solid points show mean values, error bars show +/- 1 SE, and translucent points show data from individual microcosms containing 5 mosquito larvae each.

### Tables

**Table S1. Geographic locations of field sites in from which experimental populations were sourced in California, USA. Listed from north to south.**

| region | population (county) | coordinates | distance to nearest site (km) |
|---|---|---|---|
| *northern* | Marin | 38.130, -122.544 | 71 |
| | Alameda | 37.724, -121.911 | 44 |
| | San Mateo | 37.411, -122.237 | 44 |
| *southern* | Santa Barbara | 34.543, -119.822 | 124 |
| | Los Angeles | 34.093, -118.592 | 124 |
| | San Diego | 32.681, -116.819 | 225 |

**Table S2. Summary of January daily temperature estimates for all population source locations from 1988 – 2023 PRISM estimates.** Tmax = daily maximum temperature, Tmin = daily minimum temperature. The second to last column contextualizes the rarity of the 24ºC daytime experimental heatwave temperature at the field sites. The rightmost column shows the mean daily minimum field temperatures associated with daytime temperatures similar to the heatwave treatment daytime temperatures.

| Region | Population | mean Tmax | mean Tmin | 97.5th percentile Tmax | highest recorded Tmax | 24ºC Tmax percent rank | mean Tmin for 22ºC < Tmax < 26ºC |
|---|---|---|---|---|---|---|---|
| *northern* | Marin | 13.5ºC | 4ºC | 19.5ºC | 26.9ºC | 99.9% | 5.4ºC |
| | Alameda | 13.9ºC | 3.8ºC | 19.9ºC | 24ºC | 100.0% | 5.4ºC |
| | San Mateo | 15.3ºC | 5.5ºC | 21.8ºC | 25.4ºC | 99.9% | 8.6ºC |
| *southern* | Santa Barbara | 17.8ºC | 4.7ºC | 25.9ºC | 30.2ºC | 92.5% | 5.7ºC |
| | Los Angeles | 18.7ºC | 9.5ºC | 28.4ºC | 32.3ºC | 83.8% | 12.6ºC |
| | San Diego | 18.7ºC | 5.2ºC | 27.6ºC | 30.6ºC | 86.8% | 6.6ºC |
| **Overall** | | **16.3ºC** | **5.5**ºC | **23.8**ºC | **28.2ºC** | **93.8%** | **7.4ºC** |

**Table S3. Final counts of wells in the experiment, excluding wells lost due to human error, and grouped by parasite exposure treatment, heatwave treatment, and geography.** The Marin and Los Angeles county populations (in gray) were excluded from the main analyses due to low infection rates.

| **parasite exposure** | **region** | **heatwave treatment** | **number of wells per population (5 mosquito larvae per well)** | | |
|---|---|---|---|---|---|
| *present* | *northern* | | *Alameda* | *San Mateo* | *Marin* |
| | | Control | 7 | 10 | 4 |
| | | Early | 8 | 11 | 7 |
| | | Late | 9 | 9 | 6 |
| *present* | *southern* | | *San Diego* | *Santa Barbara* | *Los Angeles* |
| | | Control | 12 | 14 | 10 |
| | | Early | 8 | 10 | 9 |
| | | Late | 12 | 7 | 8 |
| *absent* | *northern* | | *Alameda* | *San Mateo* | *Marin* |
| | | Control | 5 | 8 | 4 |
| | | Early | 5 | 8 | 4 |
| | | Late | 3 | 8 | 4 |
| *absent* | *southern* | | *San Diego* | *Santa Barbara* | *Los Angeles* |
| | | Control | 8 | 6 | 10 |
| | | Early | 8 | 8 | 9 |
| | | Late | 9 | 5 | 9 |

**Table S4. Parasite attack (encystment) model results estimating effects of heatwave treatment, checkpoint, and geographic region. Significant predictors ($p < 0.05$) are highlighted with bold text.**

| *Predictors* | *Estimate* | *Standard Error* | *Z-statistic* | *p-value* |
|---|---|---|---|---|
| **(Intercept)** | **0.84** | **0.21** | **4.08** | **$4.53 \times 10^{-5}$** |
| Early heatwave | -0.11 | 0.21 | -0.53 | 0.6 |
| **Check 2** | **-0.48** | **0.12** | **-4.02** | **$5.79 \times 10^{-5}$** |
| Southern region | -0.08 | 0.21 | 0.4 | 0.69 |
| Early heatwave x check 2 | -0.13 | 0.21 | -0.63 | 0.53 |
| Early heatwave x southern region | -0.16 | 0.21 | -0.79 | 0.43 |
| **Check 2 x southern region** | **0.54** | **0.12** | **4.51** | **$6.64 \times 10^{-6}$** |
| Early heatwave x check 2 x southern region | 0.11 | 0.21 | 0.52 | 0.61 |

| *Random effects* | *intercept variance* | *standard deviation* |
|---|---|---|
| well: population | $1.4 \times 10^{-8}$ | $1.18 \times 10^{-4}$ |
| population | 0.11 | 0.33 |

AIC: 724.7

Table note: Proportion larvae with cysts ~ early heatwave * checkpoint * geographic region + (1|population/well); modeled as a binomial distribution using glmmTMB.

**Table S5. Encystment regional model emmeans pairwise contrasts for significant predictors of interest (Table S4), with Bonferroni-adjusted p-values.**

| Interaction | Region | Check | Estimate | SE | t-ratio | p-value |
|---|---|---|---|---|---|---|
| checkpoint x region | northern, southern | 1 | 1.52 | 0.47 | 3.23 | $1.2 \times 10^{-3}$* |
| | northern, southern | 2 | -0.86 | 0.43 | -2.02 | 0.04* |
| | northern | 1, 2 | 2.29 | 0.34 | 6.84 | $<1.0 \times 10^{-4}$* |
| | southern | 1, 2 | -0.1 | 0.26 | -0.35 | 0.73 |

**Table S6. Six population parasite attack (encystment) model results estimating effects of heatwave treatment, checkpoint, and population ID. Significant predictors ($p < 0.05$) are highlighted with bold text.**

| *Predictors* | *estimate* | *standard error* | *Z-statistic* | *p-value* |
|---|---|---|---|---|
| **(Intercept)** | **0.48** | **0.11** | **4.46** | **$8.21 \times 10^{-6}$** |
| early heatwave | -0.17 | 0.18 | -0.95 | 0.34 |
| check 2 | -0.17 | 0.10 | -1.63 | 0.10 |
| San Mateo | -0.26 | 0.23 | -1.11 | 0.27 |
| **Marin** | **-0.99** | **0.28** | **-3.56** | **$3.7 \times 10^{-4}$** |
| San Diego | 0.17 | 0.19 | 0.89 | 0.37 |
| **Santa Barbara** | **0.44** | **0.21** | **2.08** | **0.04** |
| **Los Angeles** | **-0.66** | **0.21** | **-3.22** | **$1.28 \times 10^{-3}$** |
| early heatwave x check2 | -0.28 | 0.18 | -1.62 | 0.11 |
| **early heatwave x San Mateo** | **0.77** | **0.39** | **1.99** | **$4.63 \times 10^{-2}$** |
| early heatwave x Marin | 0.06 | 0.44 | 0.14 | 0.89 |
| early heatwave x San Diego | -0.09 | 0.36 | -0.25 | 0.80 |
| early heatwave x Santa Barbara | -0.12 | 0.35 | -0.34 | 0.73 |
| early heatwave x Los Angeles | -0.14 | 0.36 | -0.39 | 0.70 |
| **check 2 x San Mateo** | **-1.17** | **0.23** | **-5.10** | **$3.34 \times 10^{-7}$** |
| **check 2 x Marin** | **0.66** | **0.27** | **2.45** | **0.01** |
| check 2 x San Diego | 0.22 | 0.18 | 1.18 | 0.24 |
| | 0.23 | 0.21 | 1.11 | 0.27 |

| | | | | |
|---|---|---|---|---|
| check 2 x Santa Barbara | | | | |
| **check 2 x Los Angeles** | **0.62** | **0.20** | **3.09** | **$2.02 \times 10^{-3}$** |
| early heatwave x check 2 x San Mateo | 0.47 | 0.38 | 1.25 | 0.21 |
| **early heatwave x check 2 x Marin** | **-1.07** | **0.43** | **-2.48** | **0.01** |
| early heatwave x check 2 x San Diego | -0.14 | 0.35 | -0.41 | 0.68 |
| early heatwave x check 2 x Santa Barbara | 0.58 | 0.34 | 1.71 | 0.09 |
| early heatwave x check 2 x Los Angeles | 0.58 | 0.36 | 1.64 | 0.10 |

| *Random effects* | *intercept variance* | *standard deviation* |
|---|---|---|
| well | 0.04 | 0.2 |

AIC: 1022.1

Table notes: For all six populations, proportion larvae with cysts ~ early * check * population + (1|well) modeled as a betabinomial distribution with glmmTMB.

**Table S7. Six population parasite attack (encystment) model emmeans pairwise contrasts for significant predictors of interest (Table S6), with Bonferroni-adjusted p-values.** Populations excluded from the main analyses due to low infection are italicized.

| | Contrasts | | | | | | |
|---|---|---|---|---|---|---|---|
| **Interaction** | **Population (Italics = low infection rate)** | **Treatment** | **Check** | **Estimate** | **SE** | **Z ratio** | **p-value** |
| checkpoint x population | *Marin, Los Angeles* | - | 1 | 0.56 | 0.42 | 1.32 | 1.0 |
| | *Marin*, Alameda | - | 1 | 2.91 | 0.56 | 5.24 | <1 x $10^{-4}$* |
| | *Marin*, San Mateo | - | 1 | 2.14 | 0.45 | 4.74 | <1 x $10^{-4}$* |
| | *Marin*, Santa Barbara | - | 1 | 0.94 | 0.40 | -2.32 | 0.32 |
| | *Marin,* San Diego | - | 1 | -1.01 | 0.42 | -2.54 | 0.17 |
| | *Los Angeles*, Alameda | - | 1 | 3.47 | 0.53 | 6.54 | <1 x $10^{-4}$* |
| | *Los Angeles*, San Mateo | - | 1 | 2.70 | 0.42 | 6.44 | <1 x $10^{-4}$* |
| | *Los Angeles*, Santa Barbara | - | 1 | 1.49 | 0.37 | 4.08 | 9 x $10^{-4}$* |
| | *Los Angeles*, San Diego | - | 1 | 1.62 | 0.38 | 4.24 | 4 x $10^{-4}$* |
| | *Marin, Los Angeles* | - | 2 | -1.0 | 0.44 | -2.27 | 0.36 |
| | *Marin*, Alameda | - | 2 | 0.62 | 0.44 | 1.41 | 1.0 |
| | *Marin*, San Mateo | - | 2 | 0.03 | 0.44 | 0.06 | 1.0 |
| | *Marin*, Santa Barbara | - | 2 | -1.71 | 0.44 | -3.92 | 1.6 x $10^{-3}$* |
| | *Marin,* San Diego | - | 2 | -1.03 | 0.44 | -2.37 | 0.28 |
| | *Los Angeles*, Alameda | - | 2 | -0.38 | 0.37 | -1.03 | 1.0 |
| | *Los Angeles*, San Mateo | - | 2 | -0.97 | 0.37 | -2.66 | 0.12 |
| | *Los Angeles*, Santa Barbara | - | 2 | 0.71 | 0.36 | 1.96 | 0.77 |
| | *Los Angeles*, San Diego | - | 2 | 0.03 | 0.36 | 0.09 | 1.0 |
| | *Marin* | - | 1, 2 | 0.40 | 0.48 | 0.84 | 0.4 |
| | *Los Angeles* | - | 1, 2 | -1.15 | 0.38 | -3.07 | 2.4 x $10^{-3}$* |
| | Alameda | - | 1, 2 | 2.7 | 0.53 | 5.14 | <1.0 x $10^{-4}$* |
| | San Mateo | - | 1, 2 | 2.52 | 0.41 | 6.16 | <1.0 x $10^{-4}$ |
| | Santa Barbara | - | 1, 2 | -0.37 | 0.35 | -1.04 | 0.3 |
| | San Diego | - | 1, 2 | 0.44 | 0.37 | 1.18 | 0.24 |
| checkpoint x population x early heatwave | Marin | control, early | 1 | -1.25 | 0.65 | -1.9 | 0.06 |

| | | | | | | |
|---|---|---|---|---|---|---|
| Marin | control, early | 2 | 1.46 | 0.73 | 2.01 | 0.04* |
| Marin | control | 1, 2 | -0.99 | 0.61 | -1.61 | 0.11 |
| Marin | early | 1,2 | 1.72 | 0.75 | 2.3 | 0.02* |

**Table S8. Infection model results estimating interactive effects of heatwave treatment, checkpoint, and geographic region. Significant predictors ($p < 0.05$) are highlighted with bold text.**

| *Predictors* | *Estimate* | *Standard Error* | *Z-statistic* | *p-value* |
|---|---|---|---|---|
| **(Intercept)** | **-1.23** | **0.24** | **-5.20** | **$1.95 \times 10^{-7}$** |
| **Early heatwave** | **-2.48** | **0.41** | **-6.02** | **$1.75 \times 10^{-9}$** |
| Late heatwave | -0.42 | 0.33 | -1.28 | 0.20 |
| **Check 3** | **0.57** | **0.16** | **3.69** | **$2.21 \times 10^{-4}$** |
| **Check 4** | **0.57** | **0.16** | **3.68** | **$2.38 \times 10^{-4}$** |
| **Check 5** | **-0.44** | **0.16** | **-2.69** | **0.01** |
| **Southern region** | **-1.08** | **0.24** | **-4.58** | **$4.58 \times 10^{-6}$** |
| Early heatwave x check 3 | -0.13 | 0.38 | -0.33 | 0.74 |
| Late heatwave x check 3 | 0.23 | 0.23 | 1.00 | 0.32 |
| Early heatwave x check 4 | -0.58 | 0.51 | -1.14 | 0.25 |
| **Late heatwave x check 4** | **-1.00** | **0.27** | **-3.63** | **$2.85 \times 10^{-4}$** |
| Early heatwave x check 5 | 0.08 | 0.52 | 0.16 | 0.88 |
| Late heatwave x check 5 | -0.40 | 0.28 | -1.42 | 0.16 |
| Early heatwave x southern region | 0.12 | 0.41 | 0.30 | 0.77 |
| Late heatwave x southern region | 0.06 | 0.33 | 0.17 | 0.86 |
| Southern region x check 3 | -0.11 | 0.16 | -0.69 | 0.49 |
| Southern region x check 4 | -0.17 | 0.16 | -1.09 | 0.28 |
| **Southern region x check 5** | **0.44** | **0.16** | **2.66** | **0.01** |
| Early heatwave x check 3 x southern region | 0.59 | 0.38 | 1.53 | 0.12 |
| Late heatwave x check 3 x southern region | 0.40 | 0.23 | 1.73 | 0.08 |
| Early heatwave x check 4 x southern region | -0.37 | 0.51 | -0.74 | 0.46 |
| Late heatwave x check 4 x southern region | -0.30 | 0.27 | -1.11 | 0.27 |
| Early heatwave x check 5 x southern region | -0.63 | 0.52 | -1.21 | 0.23 |
| Late heatwave x check 5 x southern region | -0.49 | 0.28 | -1.73 | 0.08 |

| *Random effects* | *intercept variance* | *standard deviation* |
|---|---|---|
| well x population | 1.58 | 1.26 |
| population | 0.02 | 0.15 |

AIC: 990

Table notes: Proportion infected larvae ~ treatment * check * region + (1|population/well), modeled as a binomial distribution with lme4.

**Table S9. Infection regional model emmeans pairwise contrasts for significant predictors of interest (Table S8), with Bonferroni-adjusted p-values. Significant p-values < 0.05 are indicated with asterisks.**

| | Contrasts | | | | | | |
|---|---|---|---|---|---|---|---|
| **Interaction** | **Region** | **Treatment** | **Check** | **Estimate** | **SE** | **Z ratio** | **p-value** |
| late heatwave x check 4 | - | control, late | 4 | -1.42 | 0.44 | -3.23 | $2.6 \times 10^{-3}$* |
| | - | late | 3, 4 | -1.23 | 0.33 | -3.75 | $5.0 \times 10^{-4}$* |
| region x check 5 | northern | - | 4, 5 | -1.06 | 0.24 | -4.41 | $<1.0 \times 10^{-4}$* |
| | southern | - | 4, 5 | -0.14 | 0.59 | -0.23 | 1.0 |
| | northern, southern | - | 5 | 1.91 | 0.52 | 3.65 | $3.0 \times 10^{-4}$* |

**Table S10. Immune melanization response model results estimating interactive effects of heatwave treatment, checkpoint, and geographic region. Significant predictors (p < 0.05) are highlighted with bold text.**

| *predictors* | *estimate* | *standard error* | *Z-statistic* | *p-value* |
|---|---|---|---|---|
| (Intercept) | -0.31 | 0.16 | -1.93 | 0.05 |
| early heatwave | -0.02 | 0.24 | -0.10 | 0.92 |
| late heatwave | -0.11 | 0.24 | -0.44 | 0.66 |
| check 3 | 0.18 | 0.16 | 1.16 | 0.25 |
| check 4 | -0.20 | 0.17 | -1.19 | 0.23 |
| check 5 | 0.02 | 0.24 | 0.07 | 0.94 |
| southern region | 0.31 | 0.16 | 1.96 | 0.05 |
| early heatwave x check 3 | -0.41 | 0.24 | -1.74 | 0.08 |
| late heatwave x check 3 | -0.02 | 0.24 | -0.07 | 0.94 |
| early heatwave x check 4 | -0.40 | 0.25 | -1.59 | 0.11 |
| **late heatwave x check 4** | 0.92 | 0.25 | 3.60 | **$3.16 \times 10^{-4}$** |
| early heatwave x check 5 | 0.10 | 0.39 | 0.25 | 0.80 |
| late heatwave x check 5 | -0.81 | 0.42 | -1.94 | 0.05 |
| **early heatwave x southern region** | -0.79 | 0.24 | -3.30 | **$9.72 \times 10^{-4}$** |
| late heatwave x southern region | -0.03 | 0.24 | -0.11 | 0.91 |
| check 3 x southern region | -0.30 | 0.16 | -1.90 | 0.06 |
| check 4 x southern region | 0.31 | 0.17 | 1.81 | 0.07 |
| check 5 x southern region | 0.31 | 0.24 | 1.31 | 0.19 |
| **early heatwave x** | 0.73 | 0.24 | 3.08 | **$2.06 \times 10^{-3}$** |

| | | | | |
|---|---|---|---|---|
| **check 3 x southern region** | | | | |
| late heatwave x check 3 x southern region | -0.01 | 0.24 | -0.03 | 0.98 |
| early heatwave x check 4 x southern region | -0.17 | 0.25 | -0.66 | 0.51 |
| **late heatwave x check 4 x southern region** | -0.53 | 0.25 | -2.10 | **0.04** |
| early heatwave x check 5 x southern region | -0.22 | 0.39 | -0.57 | 0.57 |
| late heatwave x check 5 x southern region | 0.52 | 0.41 | 1.26 | 0.21 |

| *Random effects* | *intercept variance* | *standard deviation* |
|---|---|---|
| well | 0.55 | 0.74 |

AIC: 1146.9

Table notes: Proportion larvae melanizing ~ treatment * check * region + (1 | well), modeled as a binomial distribution with glmer.

**Table S11. Immune melanization regional model emmeans pairwise contrasts for significant predictors of interest (Table S10), with Bonferroni-adjusted p-values.**

| | Contrasts | | | | | | |
|---|---|---|---|---|---|---|---|
| **Interaction** | **Region** | **Treatment** | **Check** | **Estimate** | **SE** | **Z ratio** | **p-value** |
| early heatwave x checkpoint x region | northern | early, control | 2 | -1.82 | 0.45 | -4.0 | $2 \times 10^{-4}$* |
| | southern | early, control | 2 | 0.44 | 0.38 | 1.15 | 0.75 |
| | northern, southern | early | 2 | 2.26 | 0.44 | 5.13 | $<1 \times 10^{-4}$* |
| late heatwave x checkpoint x region | northern | late, control | 4 | -1.37 | 0.49 | -2.78 | 0.02* |
| | southern | late, control | 4 | -0.25 | 0.4 | -0.63 | 1 |
| | northern, southern | late | 4 | -0.12 | 0.45 | -0.26 | 0.8 |

**Table S12. Host pupation hazard discrete time-to-event model results estimating effects of checkpoint, parasite exposure, heatwave treatment, and geographic region. Significant predictors ($p < 0.05$) are highlighted with bold text.**

| *Predictors* | *Estimate* | *Standard Error* | *Z-statistic* | *p-value* |
|---|---|---|---|---|
| **(Intercept)** | **-3.09** | **0.28** | **-10.89** | **$1.30 \times 10^{-27}$** |
| **Check 5** | **2.75** | **0.24** | **11.25** | **$2.42 \times 10^{-29}$** |
| **Check 6** | **3.93** | **0.28** | **13.90** | **$5.94 \times 10^{-44}$** |
| **Check 7+** | **4.02** | **0.50** | **8.09** | **$5.76 \times 10^{-16}$** |
| **Parasite absent** | **1.41** | **0.27** | **5.33** | **$9.61 \times 10^{-8}$** |
| **Early heatwave** | **1.09** | **0.29** | **3.76** | **$1.68 \times 10^{-4}$** |
| **Late heatwave** | **0.95** | **0.30** | **3.23** | **$1.22 \times 10^{-3}$** |
| Southern region | -0.09 | 0.17 | -0.53 | 0.60 |
| Parasite absent x early heatwave | -0.48 | 0.28 | -1.70 | 0.09 |
| Parasite absent x late heatwave | -0.30 | 0.29 | -1.06 | 0.29 |
| Parasite absent x southern region | -0.08 | 0.20 | -0.40 | 0.69 |
| Early heatwave x southern region | -0.02 | 0.18 | -0.09 | 0.93 |
| Late heatwave x southern region | 0.05 | 0.18 | 0.27 | 0.79 |
| Check 5 x early heatwave | -0.20 | 0.26 | -0.78 | 0.43 |
| Check 6 x early heatwave | -0.63 | 0.37 | -1.71 | 0.09 |
| Check 7+ x early heatwave | -2.21 | 1.17 | -1.89 | 0.06 |
| Check 5 x late heatwave | -0.51 | 0.26 | -1.93 | 0.05 |
| **Check 6 x late heatwave** | **-1.22** | **0.33** | **-3.65** | **$2.64 \times 10^{-4}$** |
| **Check 7+ x late heatwave** | **-2.02** | **0.63** | **-3.23** | **$1.26 \times 10^{-3}$** |
| **Check 5 x parasite absent** | **-0.85** | **0.20** | **-4.21** | **$2.61 \times 10^{-5}$** |
| Check 6 x parasite absent | -0.41 | 0.30 | -1.39 | 0.16 |

| | | | | |
|---|---|---|---|---|
| **Check 7+ x parasite absent** | **-1.91** | **0.75** | **-2.53** | **0.01** |
| Parasite absent x early heatwave x southern region | -0.01 | 0.27 | -0.05 | 0.96 |
| Parasite absent x late heatwave x southern region | 0.16 | 0.28 | 0.57 | 0.57 |

| *Random effects* | *Intercept variance* | *standard deviation* |
|---|---|---|
| population | 0.05 | 0.22 |
| well:population | 0.29 | 0.54 |

AIC: 1911.0

Table note: pupation ~ checkpoint + parasite * heatwave * region + (heatwave * checkpoint) + (parasite * checkpoint) + (1 | population/well), modeled as a binomial distribution with log-log link using lme4.

**Table S13. Pupation hazard regional emmeans pairwise contrasts for significant predictors of interest (Table S12). Bonferroni-adjusted p-values <0.05 are marked with asterisks.**

| Interaction | Checkpoint | Treatment | Estimate | SE | Z ratio | p-value |
|---|---|---|---|---|---|---|
| Checkpoint x heatwave | 4 | Early heatwave, control | 0.85 | 0.24 | 3.56 | $7 \times 10^{-4}$* |
| | 4 | Late heatwave, control | 0.80 | 0.24 | 3.29 | $2 \times 10^{-3}$* |
| | 5 | Early heatwave, control | 0.65 | 0.16 | 4.10 | $<1 \times 10^{-4}$ |
| | 5 | Late heatwave, control | 0.29 | 0.16 | 1.79 | 0.15 |
| | 6 | Early heatwave, control | 0.24 | 0.30 | 0.82 | 0.83 |
| | 6 | Late heatwave, control | -0.40 | 0.25 | -1.58 | 0.23 |
| | 7+ | Early heatwave, control | -16.26 | 2480.0 | -0.007 | 1.0 |
| | 7+ | Late heatwave, control | -2.13 | 0.76 | -2.81 | 0.01* |
| Checkpoint x parasite exposure | 4 | Parasite present, absent | -1.15 | 0.19 | -6.06 | $<1 \times 10^{-4}$* |
| | 5 | Parasite present, absent | -0.29 | 0.13 | -2.24 | 0.03* |
| | 6 | Parasite present, absent | -0.71 | 0.24 | -2.94 | $3.3 \times 10^{-3}$* |
| | 7+ | Parasite present, absent | 1.24 | 0.89 | 1.40 | 0.16 |

**Table S14. Host survival to pupation model results estimating effects of parasite exposure, heatwave treatment, and geographic region. Significant predictors ($p < 0.05$) are highlighted with bold text.**

| *Predictors* | *Estimate* | *Standard Error* | *Z-statistic* | *p-value* |
|---|---|---|---|---|
| **(Intercept)** | **0.85** | **0.20** | **4.34** | **$1.41 \times 10^{-5}$** |
| **Early heatwave** | **1.35** | **0.29** | **4.61** | **$4.09 \times 10^{-6}$** |
| Late heatwave | 0.19 | 0.23 | 0.80 | 0.42 |
| **Parasite absent** | **2.44** | **0.49** | **4.98** | **$6.51 \times 10^{-7}$** |
| Southern region | 0.27 | 0.20 | 1.40 | 0.16 |
| **Early heatwave x parasite absent** | **-1.50** | **0.69** | **-2.18** | **0.03** |
| Late heatwave x parasite absent | -0.29 | 0.70 | -0.42 | 0.67 |
| Early heatwave x southern region | 0.00 | 0.29 | 0.01 | 0.99 |
| Late heatwave x southern region | 0.05 | 0.23 | 0.23 | 0.82 |
| Parasite absent x southern region | -0.42 | 0.49 | -0.86 | 0.39 |
| Early heatwave x parasite absent x southern region | 0.27 | 0.69 | 0.39 | 0.69 |
| Late heatwave x parasite absent x southern region | 0.07 | 0.70 | 0.10 | 0.92 |

| *Random effects* | *intercept variance* | *standard deviation* |
|---|---|---|
| population | 0.06 | 0.24 |

AIC: 425

Table note: Survival to pupation ~ heatwave treatment * parasite exposure * region + (1 | population), modeled as a binomial distribution using lme4.

**Table S15. Host survival to pupation regional model emmeans pairwise contrasts for significant predictors of interest (Table S14). Bonferroni-adjusted p-values <0.05 are marked with asterisks.**

| | Contrasts | | | | | | |
|---|---|---|---|---|---|---|---|
| **Interaction** | **Region** | **Heatwave treatment** | **parasite** | **Estimate** | **SE** | **Z ratio** | **p-value** |
| treatment x parasite exposure | - | early, control | present | -1.35 | 0.29 | -4.61 | $<1 \times 10^{-4}$* |
| | - | late, control | present | 0.19 | 0.23 | 0.81 | 0.84 |
| | - | early, control | absent | -0.15 | 0.62 | -0.24 | 1.0 |
| | - | late, control | absent | -0.11 | 0.66 | -0.17 | 1.0 |
| | - | early | present, absent | -0.93 | 0.49 | -1.92 | 0.05 |
| | - | late | present, absent | -2.14 | 0.49 | -4.32 | $<1 \times 10^{-4}$* |
| | - | control | present, absent | -2.44 | 0.49 | -4.98 | $<1 \times 10^{-4}$* |

**Table S16. Parasite attack (encystment) population robustness check model results estimating effects of three-way interactions between heatwave treatment, checkpoint, and population. Significant predictors ($p < 0.05$) are highlighted with bold text.**

| *Predictors* | *Estimate* | *Standard Error* | *Z-statistic* | *p-value* |
|---|---|---|---|---|
| **(Intercept)** | 0.87 | 0.13 | 6.89 | $5.66 \times 10^{-12}$ |
| Early heatwave | -0.15 | 0.21 | -0.69 | 0.49 |
| **Check 2** | **-0.48** | **0.13** | **-3.83** | **$1.3 \times 10^{-4}$** |
| **San Mateo** | **-0.66** | **0.22** | **-3.08** | **$2.04 \times 10^{-3}$** |
| San Diego | -0.24 | 0.18 | -1.29 | 0.20 |
| Santa Barbara | 0.03 | 0.20 | 0.16 | 0.88 |
| Early heatwave x check 2 | -0.16 | 0.21 | -0.75 | 0.45 |
| **Early heatwave x San Mateo** | **0.74** | **0.36** | **2.05** | **0.04** |
| Early heatwave x San Diego | -0.11 | 0.34 | -0.31 | 0.76 |
| Early heatwave x Santa Barbara | -0.15 | 0.33 | -0.44 | 0.66 |
| **Check 2 x San Mateo** | **-0.83** | **0.22** | **-3.86** | **$1.12 \times 10^{-4}$** |
| **Check 2 x San Diego** | **0.53** | **0.18** | **2.90** | **$3.7 \times 10^{-3}$** |
| **Check 2 x Santa Barbara** | **0.54** | **0.20** | **2.68** | **$7.44 \times 10^{-3}$** |
| Early heatwave x check 2 x San Mateo | 0.35 | 0.36 | 0.96 | 0.34 |
| Early heatwave x check 2 x San Diego | -0.27 | 0.34 | -0.77 | 0.44 |
| Early heatwave x check 2 x Santa Barbara | 0.45 | 0.33 | 1.36 | 0.18 |

| *Random effects* | *intercept variance* | *standard deviation* |
|---|---|---|
| well | $8.54 \times 10^{-9}$ | $9.24 \times 10^{-5}$ |

AIC: 718.3

Table notes: Proportion larvae with cysts ~ early * check * population + (1|well), modeled as a betabinomial distribution with glmmTMB.

**Table S17. Encystment population robustness check emmeans pairwise contrasts for significant predictors of interest in the population-level three-way interaction model (Table S16). Bonferroni-adjusted p-values < 0.05 are marked with asterisks.**

| Predictors | Population | Treatment | Check | Estimate | SE | t-ratio | p-value |
|---|---|---|---|---|---|---|---|
| check * population | Alameda, San Mateo | | 1 | 0.76 | 0.56 | 1.37 | 1.0 |
| | Alameda, San Diego | | 1 | 1.82 | 0.53 | 3.42 | $4.5 \times 10^{-3}$* |
| | Alameda, Santa Barbara | | 1 | 1.93 | 0.52 | 3.71 | $1.6 \times 10^{-3}$* |
| | San Mateo, San Diego | | 1 | 1.06 | 0.42 | 2.61 | 0.08 |
| | San Mateo, Santa Babara | | 1 | 1.17 | 0.41 | 2.87 | 0.03* |
| | San Diego, Santa Barbara | | 1 | 0.12 | 0.37 | 0.31 | 1.0 |
| | Alameda | | 1, 2 | 2.13 | 0.54 | 3.94 | $1 \times 10^{-4}$* |
| | San Mateo | | 1, 2 | 2.44 | 0.42 | 5.88 | $<1 \times 10^{-4}$* |
| | San Diego | | 1, 2 | 0.32 | 0.38 | 0.84 | 0.4 |
| | Santa Barbara | | 1, 2 | -0.41 | 0.36 | -1.12 | 0.26 |
| early heatwave x population | Alameda | control, early | - | 0.64 | 0.54 | 1.18 | 0.24 |
| | San Mateo | control, early | - | -0.59 | 0.41 | -1.43 | 0.15 |
| | San Diego | control, early | - | 0.26 | 0.38 | 0.67 | 0.5 |
| | Santa Barbara | control, early | - | 0.3 | 0.36 | 0.82 | 0.41 |

**Table S18. Simplified encystment robustness check results estimating interactive effects of heatwave treatment and checkpoint and additive effects of population during the first two checkpoints. Significant predictors ($p < 0.05$) are highlighted with bold text.**

| *Predictors* | *Estimate* | *Standard Error* | *Z-statistic* | *p-value* |
|---|---|---|---|---|
| **(Intercept)** | **0.81** | **0.12** | **6.73** | **$1.73 \times 10^{-11}$** |
| Early heatwave | -0.13 | 0.21 | -0.64 | 0.52 |
| **Check 2** | **-0.37** | **0.12** | **-3.20** | **$1.36 \times 10^{-3}$** |
| **San Mateo** | **-0.38** | **0.16** | **-2.39** | **0.02** |
| San Diego | -0.22 | 0.16 | -1.42 | 0.16 |
| Santa Barbara | 0.06 | 0.17 | 0.38 | 0.71 |
| Early heatwave x check 2 | -0.19 | 0.21 | -0.90 | 0.37 |

| *Random effects* | *intercept variance* | *standard deviation* |
|---|---|---|
| well | $3.63 \times 10^{-9}$ | $6.03 \times 10^{-5}$ |

AIC: 746.9

Table notes: Proportion larvae with cysts ~ early * check * population + (1|well), modeled as a betabinomial distribution with glmmTMB.

**Table S19. Simplified encystment robustness check emmeans pairwise contrasts for significant predictors of interest in the population-level two-way interaction model (Table S18), with Bonferroni-adjusted p-values.**

| Predictors | Population | Check | Estimate | SE | t-ratio | p-value |
|---|---|---|---|---|---|---|
| check * population | Alameda, San Mateo | - | 0.93 | 0.29 | 3.19 | $9.0 \times 10^{-3}$* |
| | Alameda, San Diego | - | 0.77 | 0.29 | 2.67 | $4.97 \times 10^{-2}$* |
| | Alameda, Santa Barbara | - | 0.48 | 0.30 | 1.62 | 0.64 |
| | San Mateo, San Diego | - | -0.16 | 0.25 | -0.64 | 1.0 |
| | San Mateo, Santa Babara | - | -0.45 | 0.26 | -1.7 | 0.54 |
| | San Diego, Santa Barbara | - | -0.29 | 0.26 | -1.11 | 1.0 |

**Table S20. Infection population robustness check results estimating interactive effects of heatwave treatment, checkpoint, and population. Significant predictors ($p < 0.05$) are highlighted with bold text.**

| *Predictors* | *Estimate* | *Standard Error* | *Z-statistic* | *p-value* |
|---|---|---|---|---|
| **(Intercept)** | **-1.29** | **0.19** | **-6.86** | **$6.91 \times 10^{-12}$** |
| **Early heatwave** | **-1.88** | **0.31** | **-6.09** | **$1.1 \times 10^{-9}$** |
| Lat heatwave | -0.24 | 0.27 | -0.91 | $3.6 \times 10^{-1}$ |
| **check3** | **0.53** | **0.14** | **3.75** | **$1.77 \times 10^{-4}$** |
| **check4** | **0.46** | **0.14** | **3.20** | **$1.39 \times 10^{-3}$** |
| **check5** | **-0.40** | **0.15** | **-2.63** | **$8.43 \times 10^{-3}$** |
| San Mateo | 0.36 | 0.26 | 1.36 | 0.17 |
| **San Diego** | **-1.04** | **0.27** | **-3.89** | **$1.0 \times 10^{-4}$** |
| **Santa Barbara** | **-0.67** | **0.26** | **-2.63** | **$8.6 \times 10^{-3}$** |
| Early heatwave x check 3 | -0.24 | 0.28 | -0.85 | 0.39 |
| Late heatwave x check 3 | 0.21 | 0.21 | 1.00 | 0.32 |
| Early heatwave x check 4 | -0.27 | 0.29 | -0.92 | 0.36 |
| **Late heatwave x check 4** | **-0.75** | **0.23** | **-3.23** | **1.24E-03** |
| Early heatwave x check 5 | 0.11 | 0.31 | 0.36 | 0.72 |
| Late heatwave x check 5 | -0.39 | 0.24 | -1.62 | 0.11 |
| Early heatwave x San Mateo | 0.03 | 0.38 | 0.07 | 0.94 |
| Late heatwave x San Mateo | 0.24 | 0.36 | 0.67 | 0.5 |
| Early heatwave x San Diego | 0.44 | 0.42 | 1.05 | 0.29 |
| Late heatwave x San Diego | -0.42 | 0.37 | -1.14 | 0.25 |
| Early heatwave x Santa Barbara | -0.26 | 0.41 | -0.62 | 0.53 |
| Late heatwave x Santa Barbara | 0.37 | 0.38 | 0.98 | 0.33 |
| San Mateo x check 3 | 0.06 | 0.20 | 0.29 | 0.77 |
| San Mateo x check 4 | 0.28 | 0.21 | 1.37 | 0.17 |
| San Mateo x check 5 | -0.35 | 0.22 | -1.60 | 0.11 |
| San Diego x check 3 | -0.19 | 0.23 | -0.84 | 0.4 |
| San Diego x check 4 | -0.21 | 0.23 | -0.90 | 0.37 |
| San Diego x check 5 | 0.26 | 0.24 | 1.05 | 0.29 |
| Santa Barbara x check 3 | 0.13 | 0.21 | 0.61 | 0.54 |
| Santa Barbara x check 4 | -0.10 | 0.22 | -0.48 | 0.63 |
| Santa Barbara x check 5 | 0.29 | 0.23 | 1.29 | 0.2 |
| Early heatwave x San Mateo x check 3 | 0.16 | 0.34 | 0.47 | 0.64 |
| Late heatwave x San Mateo x check 3 | -0.01 | 0.29 | -0.03 | 0.98 |
| Early heatwave x San Mateo x check 4 | 0.13 | 0.34 | 0.39 | 0.7 |
| Late heatwave x San Mateo x check 4 | -0.21 | 0.30 | -0.71 | 0.48 |
| Early heatwave x San Mateo x check 5 | 0.13 | 0.37 | 0.34 | 0.73 |

| | | | | |
|---|---|---|---|---|
| Late heatwave x San Mateo x check 5 | -0.11 | 0.32 | -0.34 | 0.73 |
| Early heatwave x San Diego x check 3 | 0.47 | 0.40 | 1.17 | 0.24 |
| Late heatwave x San Diego x check 3 | 0.24 | 0.33 | 0.75 | 0.45 |
| Early heatwave x San Diego x check 4 | -0.07 | 0.41 | -0.18 | 0.86 |
| Late heatwave x San Diego x check 4 | 0.28 | 0.36 | 0.80 | 0.42 |
| Early heatwave x San Diego x check 5 | -0.17 | 0.42 | -0.40 | 0.69 |
| Late heatwave x San Diego x check 5 | 0.27 | 0.36 | 0.75 | 0.46 |
| Early heatwave x Santa Barbara x check 3 | 0.04 | 0.41 | 0.11 | 0.91 |
| Late heatwave x Santa Barbara x check 3 | 0.23 | 0.31 | 0.75 | 0.45 |
| Early heatwave x Santa Barbara x check 4 | -0.27 | 0.43 | -0.63 | 0.53 |
| **Late heatwave x Santa Barbara x check 4** | **-0.70** | **0.35** | **-2.00** | **$4.5 \times 10^{-2}$** |
| Early heatwave x Santa Barbara x check 5 | -0.37 | 0.44 | -0.83 | 0.41 |
| Late heatwave x Santa Barbara x check 5 | -0.69 | 0.36 | -1.92 | $5.4 \times 10^{-2}$ |

| *Random effects* | *intercept variance* | *standard deviation* |
|---|---|---|
| well | 1.31 | 1.15 |

AIC: 1005

Table note: Proportion larvae infected ~ treatment * check *  population + (1 | well), modeled as a binomial distribution with weak priors on the fixed effects using blme.

**Table S21. Infection population robustness check emmeans pairwise contrasts for significant predictors of interest the population-level three-way interaction model (Table S20). Bonferroni-adjusted p-values <0.05 are indicated with asterisks.**

| Predictors | Population | Treatment | Check | Estimate | SE | Z ratio | p-value |
|---|---|---|---|---|---|---|---|
| late heatwave x check 4 | all | control, late | 4 | -0.99 | 0.36 | -2.77 | 0.01* |
| | all | late | 3, 4 | -1.03 | 0.29 | -3.55 | $1.2 \times 10^{-3}$* |
| late heatwave x check 4 x population | Alameda | control, late | 4 | -0.56 | 0.67 | -0.83 | 0.81 |
| | San Mateo | control, late | 4 | -0.96 | 0.56 | -1.71 | 0.17 |
| | San Diego | control, late | 4 | -1.13 | 0.64 | -1.76 | 0.16 |
| | Santa Barbara | control, late | 4 | -1.32 | 0.67 | -1.99 | 0.09 |
| | Alameda | late | 3, 4 | 0.1 | 0.46 | -0.22 | 1.0 |
| | San Mateo | late | 3, 4 | -1.01 | 0.46 | 2.23 | 0.08 |
| | San Diego | late | 3, 4 | -1.01 | 0.61 | 1.66 | 0.29 |
| | Santa Barbara | late | 3, 4 | -2.2 | 0.62 | 3.55 | $1.2 \times 10^{-3}$* |

**Table S22. Simplified infection robustness check results estimating interactive effects of heatwave treatment and checkpoint and additive effects of population. Significant predictors ($p < 0.05$) are highlighted with bold text.**

| *Predictors* | *Estimate* | *Standard Error* | *Z-statistic* | *p-value* |
|---|---|---|---|---|
| **(Intercept)** | **-1.23** | **0.22** | **-5.64** | **$1.73 \times 10^{-8}$** |
| **Early heatwave** | **-2.39** | **0.37** | **-6.46** | **$1.05 \times 10^{-10}$** |
| Late heatwave | -0.33 | 0.32 | -1.04 | 0.3 |
| **Check 3** | **0.57** | **0.15** | **3.68** | **$2.31 \times 10^{-4}$** |
| **Check 4** | **0.57** | **0.15** | **3.68** | **$2.31 \times 10^{-4}$** |
| **Check 5** | **-0.49** | **0.17** | **-2.91** | **$3.58 \times 10^{-4}$** |
| **San Mateo** | **0.66** | **0.23** | **2.87** | **$4.17 \times 10^{-4}$** |
| **San Diego** | **-1.35** | **0.25** | **-5.41** | **$6.41 \times 10^{-8}$** |
| **Santa Barbara** | **-0.64** | **0.25** | **-2.63** | **$8.65 \times 10^{-3}$** |
| Early heatwave x check 3 | -0.39 | 0.31 | -1.26 | 0.21 |
| Late heatwave x check 3 | 0.14 | 0.23 | 0.61 | 0.54 |
| Early heatwave x check 4 | -0.22 | 0.31 | -0.71 | 0.48 |
| **Late heatwave x check 4** | **-0.79** | **0.24** | **-3.36** | **$7.93 \times 10^{-4}$** |
| Early heatwave x check 5 | 0.23 | 0.35 | 0.65 | 0.51 |
| Late heatwave x check 5 | -0.36 | 0.26 | -1.38 | 0.17 |

| *Random effects* | *intercept variance* | *standard deviation* |
|---|---|---|
| well | 1.43 | 1.2 |

AIC: 982

Table note: Proportion larvae infected ~ treatment * check + population + (1 | well), modeled as a binomial distribution using glmer.

**Table S23. Simplified infection robustness check emmeans pairwise contrasts for significant predictors of interest in the population-level two-way interaction model (Table S22). Bonferroni-adjusted p-values < 0.05 are indicated with asterisks.**

| Predictors | Population | Treatment | Check | Estimate | SE | Z ratio | p-value |
|---|---|---|---|---|---|---|---|
| late heatwave x check 4 | - | control, late | 4 | -1.12 | 0.39 | -2.86 | $8.5 \times 10^{-3}$* |
| | - | late | 3, 4 | -0.93 | 0.28 | -3.35 | $2.0 \times 10^{-3}$* |
| population pairs | Alameda, San Mateo | - | - | 0.68 | 0.38 | 1.8 | 0.43 |
| | Alameda, San Diego | - | - | 2.69 | 0.41 | 6.56 | $<1 \times 10^{-4}$* |
| | Alameda, Santa Barbara | - | - | 1.98 | 0.41 | 4.89 | $<1 \times 10^{-4}$* |
| | San Mateo, San Diego | - | - | 2.01 | 0.39 | 5.12 | $<1 \times 10^{-4}$* |
| | San Mateo, Santa Barbara | - | - | 1.3 | 0.39 | 3.38 | $4.4 \times 10^{-3}$* |
| | San Diego, Santa Barbara | - | - | -0.71 | 0.4 | -1.75 | 0.48 |

**Table S24. Melanization population robustness check results estimating effects of heatwave treatment, checkpoint, and population. Significant predictors ($p < 0.05$) are highlighted with bold text.**

| *Predictors* | *Estimate* | *Standard Error* | *Z-statistic* | *p-value* |
|---|---|---|---|---|
| (Intercept) | -0.26 | 0.13 | -1.92 | 0.05 |
| Early heatwave | -0.09 | 0.20 | -0.46 | 0.65 |
| Late heatwave | -0.03 | 0.19 | -0.13 | 0.90 |
| Check 3 | 0.11 | 0.13 | 0.82 | 0.41 |
| Check 4 | -0.08 | 0.14 | -0.58 | 0.56 |
| Check 5 | -0.01 | 0.17 | -0.05 | 0.96 |
| San Mateo | -0.06 | 0.19 | -0.32 | 0.75 |
| San Diego | 0.23 | 0.19 | 1.23 | 0.22 |
| Santa Barbara | 0.06 | 0.18 | 0.36 | 0.72 |
| Early heatwave x check 3 | -0.31 | 0.19 | -1.63 | 0.10 |
| Late heatwave x check 3 | -0.03 | 0.19 | -0.18 | 0.86 |
| **Early heatwave x check 4** | **-0.44** | **0.20** | **-2.18** | **0.03** |
| **Late heatwave x check 4** | **0.58** | **0.20** | **2.89** | **$3.9 \times 10^{-3}$** |
| Early heatwave x check 5 | -0.01 | 0.27 | -0.05 | 0.96 |
| Late heatwave x check 5 | -0.41 | 0.26 | -1.55 | 0.12 |
| Early heatwave x San Mateo | 0.41 | 0.27 | 1.53 | 0.13 |
| Late heatwave x San Mateo | -0.23 | 0.27 | -0.84 | 0.40 |
| Early heatwave x San Diego | -0.21 | 0.27 | -0.78 | 0.44 |
| Late heatwave x San Diego | -0.25 | 0.25 | -0.97 | 0.33 |
| **Early heatwave x Santa Barbara** | **-0.59** | **0.27** | **-2.19** | **0.03** |
| Late heatwave x Santa Barbara | 0.27 | 0.27 | 0.97 | 0.33 |
| Check 3 x San Mateo | 0.00 | 0.19 | 0.01 | 0.99 |
| Check 4 x San Mateo | -0.18 | 0.19 | -0.92 | 0.36 |
| Check 5 x San Mateo | -0.16 | 0.25 | -0.64 | 0.52 |
| Check 3 x San Diego | -0.09 | 0.18 | -0.48 | 0.63 |
| Check 4 x San Diego | 0.22 | 0.18 | 1.20 | 0.23 |
| Check 5 x San Diego | 0.39 | 0.22 | 1.76 | 0.08 |
| Check 3 x Santa Barbara | -0.15 | 0.18 | -0.83 | 0.41 |
| Check 4 x Santa Barbara | 0.05 | 0.18 | 0.28 | 0.78 |
| Check 5 x Santa Barbara | 0.08 | 0.22 | 0.37 | 0.71 |
| Early heatwave x check 3 x San Mateo | -0.14 | 0.25 | -0.56 | 0.57 |
| Late heatwave x check 3 x San Mateo | -0.03 | 0.26 | -0.11 | 0.91 |

| | | | | |
|---|---|---|---|---|
| Early heatwave x check 4 x San Mateo | 0.23 | 0.27 | 0.87 | 0.38 |
| Late heatwave x check 4 x San Mateo | 0.21 | 0.27 | 0.76 | 0.45 |
| Early heatwave x check 5 x San Mateo | -0.10 | 0.34 | -0.28 | 0.78 |
| Late heatwave x check 5 x San Mateo | 0.04 | 0.34 | 0.12 | 0.91 |
| Early heatwave x check 3 x San Diego | 0.13 | 0.26 | 0.50 | 0.62 |
| Late heatwave x check 3 x San Diego | -0.08 | 0.24 | -0.32 | 0.75 |
| Early heatwave x check 4 x San Diego | 0.17 | 0.27 | 0.62 | 0.53 |
| Late heatwave x check 4 x San Diego | -0.21 | 0.25 | -0.85 | 0.39 |
| Early heatwave x check 5 x San Diego | 0.03 | 0.32 | 0.11 | 0.91 |
| Late heatwave x check 5 x San Diego | 0.25 | 0.30 | 0.84 | 0.40 |
| **Early heatwave x check 3 x Santa Barbara** | **0.63** | **0.26** | **2.46** | **0.01** |
| Late heatwave x check 3 x Santa Barbara | -0.07 | 0.26 | -0.26 | 0.79 |
| Early heatwave x check 4 x Santa Barbara | -0.06 | 0.27 | -0.23 | 0.82 |
| Late heatwave x check 4 x Santa Barbara | -0.10 | 0.27 | -0.37 | 0.71 |
| Early heatwave x check 5 x Santa Barbara | 0.03 | 0.32 | 0.10 | 0.92 |
| Late heatwave x check 5 x Santa Barbara | 0.02 | 0.33 | 0.05 | 0.96 |

| *Random effects* | *intercept variance* | *standard deviation* |
|---|---|---|
| well | 0.5 | 0.71 |

AIC: 1184.2

Table note: Proportion larvae melanizing ~ treatment * check * population + (1 | well), modeled as a binomial distribution with weak priors on fixed effects using blme.

**Table S25. Melanization robustness check emmeans pairwise contrasts for significant predictors of interest in the population-level three-way interaction model (Table S24). Bonferroni-adjusted p-values <0.05 are marked with asterisks.**

| | Contrasts | | | | | | |
|---|---|---|---|---|---|---|---|
| **Interaction** | **Population** | **Treatment** | **Check** | **Estimate** | **SE** | **Z ratio** | **p-value** |
| early heatwave x population x check 2 | Alameda | early, control | 2 | 1.99 | 0.66 | 3.01 | $5.3 \times 10^{-3}$* |
| | San Mateo | early, control | 2 | 1.09 | 0.46 | 2.37 | 0.04* |
| | San Diego | early, control | 2 | 0.13 | 0.47 | 0.28 | 1.0 |
| | Santa Barbara | early, control | 2 | -0.52 | 0.44 | -1.18 | 0.47 |

**Table S26. Simplified melanization robustness check results estimating interactive effects of heatwave treatment and checkpoint, and additive effects of population. Significant predictors (p < 0.05) are highlighted with bold text.**

| *Predictors* | *Estimate* | *Standard Error* | *Z-statistic* | *p-value* |
|---|---|---|---|---|
| (Intercept) | -0.18 | 0.16 | -1.16 | 0.25 |
| Early heatwave | -0.16 | 0.24 | -0.68 | 0.50 |
| Late heatwave | -0.14 | 0.23 | -0.59 | 0.56 |
| Check 3 | 0.08 | 0.15 | 0.51 | 0.61 |
| Check 4 | -0.13 | 0.15 | -0.82 | 0.41 |
| Check 5 | 0.16 | 0.20 | 0.80 | 0.42 |
| San Mateo | 0.05 | 0.16 | 0.34 | 0.73 |
| San Diego | -0.03 | 0.15 | -0.17 | 0.87 |
| Santa Barbara | -0.10 | 0.15 | -0.66 | 0.51 |
| Early heatwave x check 3 | -0.30 | 0.23 | -1.30 | 0.19 |
| Late heatwave x check 3 | -0.01 | 0.22 | -0.07 | 0.95 |
| Early heatwave x check 4 | -0.46 | 0.24 | -1.89 | 0.06 |
| **Late heatwave x check 4** | **0.74** | **0.23** | **3.19** | **$1.40 \times 10^{-3}$** |
| Early heatwave x check 5 | -0.04 | 0.37 | -0.10 | 0.92 |
| Late heatwave x check 5 | -0.65 | 0.34 | -1.93 | 0.05 |

| *Random effects* | *intercept variance* | *standard deviation* |
|---|---|---|
| well | 0.6 | 0.77 |

AIC: 1179.4

Table note: Proportion larvae melanizing ~ treatment * check + population + (1 | well), modeled as a binomial distribution with glmer.

**Table S27. Simplified melanization robustness check emmeans pairwise contrasts for the two-way interaction population-level model (Table S26). Bonferroni-adjusted p-values <0.05 are marked with asterisks.**

| Interaction | Treatment | Check | Estimate | SE | Z ratio | p-value |
|---|---|---|---|---|---|---|
| late heatwave x check 4 | late, control | 4 | 0.61 | 0.32 | -1.96 | 0.1 |
| | late | 3,4 | -0.56 | 0.25 | -2.2 | 0.17 |

**Table S28. Pupation hazard robustness check results estimating interactive effects of parasite exposure, heatwave treatment, and population, and two-way interactions of each predictor with checkpoint. Significant predictors ($p < 0.05$) are highlighted with bold text.**

| *Predictors* | *Estimate* | *Standard Error* | *Z-statistic* | *p-value* |
|---|---|---|---|---|
| **(Intercept)** | **-3.06** | **0.26** | **-11.60** | **$4.27 \times 10^{-31}$** |
| **Check 5** | **2.78** | **0.25** | **11.21** | **$3.77 \times 10^{-29}$** |
| **Check 6** | **3.94** | **0.29** | **13.81** | **$2.12 \times 10^{-43}$** |
| **Check 7+** | **4.00** | **0.50** | **8.04** | **$9.26 \times 10^{-16}$** |
| **Parasite absent** | **1.36** | **0.27** | **5.12** | **$3.02 \times 10^{-7}$** |
| **Early heatwave** | **1.05** | **0.29** | **3.60** | **$3.21 \times 10^{-4}$** |
| **Late heatwave** | **0.95** | **0.30** | **3.22** | **$1.30 \times 10^{-3}$** |
| San Mateo | 0.09 | 0.21 | 0.42 | 0.67 |
| San Diego | -0.25 | 0.20 | -1.25 | 0.21 |
| Santa Barbara | -0.04 | 0.20 | -0.22 | 0.83 |
| Parasite absent x early heatwave | -0.41 | 0.28 | -1.45 | 0.15 |
| Parasite absent x late heatwave | -0.27 | 0.29 | -0.95 | 0.34 |
| Parasite absent x San Mateo | 0.51 | 0.32 | 1.60 | 0.11 |
| Parasite absent x San Diego | -0.32 | 0.31 | -1.04 | 0.30 |
| Parasite absent x Santa Barbara | 0.38 | 0.33 | 1.16 | 0.25 |
| Early heatwave x San Mateo | 0.37 | 0.30 | 1.22 | 0.22 |
| Late heatwave x San Mateo | 0.30 | 0.31 | 0.97 | 0.33 |
| Early hatwave x San Diego | 0.02 | 0.30 | 0.07 | 0.94 |
| Late heatwave x San Diego | -0.25 | 0.28 | -0.87 | 0.39 |
| Early heatwave x Santa Barbara | 0.09 | 0.28 | 0.32 | 0.75 |
| Late heatwave x Santa Barbara | 0.53 | 0.31 | 1.71 | 0.09 |
| Check 5 x early heatwave | -0.24 | 0.26 | -0.93 | 0.35 |
| Check 6 x early heatwave | -0.66 | 0.37 | -1.79 | 0.07 |
| Check 7+ x early heatwave | -2.20 | 1.18 | -1.87 | 0.06 |
| Check 5 x late heatwave | -0.53 | 0.27 | -2.00 | 0.05 |
| **Check 6 x late heatwave** | **-1.23** | **0.34** | **-3.63** | **$2.84 \times 10^{-4}$** |
| **Check 7+ x late heatwave** | **-1.96** | **0.63** | **-3.12** | **$1.8 \times 10^{-3}$** |
| **Check 5 x parasite absent** | **-0.88** | **0.20** | **-4.28** | **$1.85 \times 10^{-5}$** |
| Check 6 x parasite absent | -0.42 | 0.30 | -1.41 | 0.16 |
| **Check 7+ x parasite absent** | **-1.88** | **0.75** | **-2.50** | **0.01** |
| Parasite absent x early heatwave x San Mateo | -0.60 | 0.45 | -1.35 | 0.18 |
| Parasite absent x late heatwave x San Mateo | -0.78 | 0.46 | -1.70 | 0.09 |
| Parasite absent x early heatwave x San Diego | 0.17 | 0.45 | 0.37 | 0.71 |
| Parasite absent x late heatwave x San Diego | 0.68 | 0.43 | 1.58 | 0.11 |

| | | | | |
|---|---|---|---|---|
| Parasite absent x early heatwave x Santa Barbara | -0.44 | 0.45 | -0.99 | 0.32 |
| Parasite absent x late heatwave x Santa Barbara | -0.69 | 0.49 | -1.42 | 0.16 |

| *Random effects* | *intercept variance* | *standard deviation* |
|---|---|---|
| well | 0.26 | 0.51 |

AIC: 1920.1

Table note: Pupation ~ checkpoint + parasite * heatwave * population + (heatwave * checkpoint) + (parasite * checkpoint) + (1 | well), modeled as a binomial distribution with log-log link using lme4.

**Table S29. Pupation hazard robustness check emmeans pairwise contrasts for the population-level three-way interaction model (Table S28). Bonferroni-adjusted p-values <0.05 are marked with asterisks.**

| Interaction | Checkpoint | Treatment | Estimate | SE | Z ratio | p-value |
|---|---|---|---|---|---|---|
| Checkpoint x heatwave | 4 | Early heatwave, control | 0.85 | 0.24 | 3.53 | $8 \times 10^{-4}$* |
| | 4 | Late heatwave, control | 0.82 | 0.25 | 3.3 | $1.9 \times 10^{-3}$* |
| | 5 | Early heatwave, control | 0.61 | 0.16 | 3.9 | $2 \times 10^{-4}$* |
| | 5 | Late heatwave, control | 0.29 | 0.16 | 1.76 | 0.16 |
| | 6 | Early heatwave, control | 0.19 | 0.3 | 0.63 | 1.0 |
| | 6 | Late heatwave, control | -0.41 | 0.26 | -1.62 | 0.21 |
| | 7+ | Early heatwave, control | -1.35 | 1.15 | -1.17 | 0.48 |
| | 7+ | Late heatwave, control | -1.14 | 0.59 | -1.95 | 0.1 |
| Checkpoint x parasite exposure | 4 | Parasite present, absent | -1.14 | 0.19 | -5.93 | $<1 \times 10^{-4}$* |
| | 5 | Parasite present, absent | -0.26 | 0.13 | -1.98 | $4.7 \times 10^{-2}$* |
| | 6 | Parasite present, absent | -0.72 | 0.25 | -2.91 | $3.7 \times 10^{-3}$* |
| | 7+ | Parasite present, absent | 0.75 | 0.73 | 1.02 | 0.31 |

**Table S30. Simplified pupation hazard robustness check results estimating two-way interactions for parasite exposure, heatwave treatment, and checkpoint, and additive effects of population. Significant predictors ($p < 0.05$) are highlighted with bold text.**

| *Predictors* | *Estimate* | *Standard Error* | *Z-statistic* | *p-value* |
|---|---|---|---|---|
| **(Intercept)** | **-3.10** | **0.26** | **-12.00** | **$3.74 \times 10^{-33}$** |
| **Check 5** | **2.75** | **0.24** | **11.24** | **$2.60 \times 10^{-29}$** |
| **Check 6** | **3.93** | **0.28** | **13.92** | **$4.68 \times 10^{-44}$** |
| **Check 7+** | **4.02** | **0.50** | **8.13** | **$4.38 \times 10^{-16}$** |
| **Parasite absent** | **1.43** | **0.26** | **5.43** | **$5.5 \times 10^{-8}$** |
| **Early heatwave** | **1.10** | **0.29** | **3.83** | **$1.30 \times 10^{-4}$** |
| **Late heatwave** | **0.97** | **0.29** | **3.30** | **$9.51 \times 10^{-4}$** |
| **San Mateo** | **0.35** | **0.09** | **3.76** | **$1.71 \times 10^{-4}$** |
| **San Diego** | **-0.32** | **0.09** | **-3.54** | **$4.01 \times 10^{-4}$** |
| Santa Barbara | 0.15 | 0.10 | 1.58 | 0.11 |
| Parasite absent x early heatwave | -0.50 | 0.28 | -1.81 | 0.07 |
| Parasite absent x late heatwave | -0.31 | 0.28 | -1.09 | 0.27 |
| Check 5 x early heatwave | -0.21 | 0.26 | -0.82 | 0.41 |
| Check 6 x early heatwave | -0.63 | 0.36 | -1.72 | 0.08 |
| Check 7+ x early heatwave | -2.20 | 1.17 | -1.88 | 0.06 |
| Check 5 x late heatwave | -0.50 | 0.26 | -1.90 | 0.06 |
| **Check 6 x late heatwave** | **-1.21** | **0.33** | **-3.62** | **$2.95 \times 10^{-4}$** |
| **Check 7+ x late heatwave** | **-1.99** | **0.62** | **-3.20** | **$1.4 \times 10^{-3}$** |
| **Check 5 x parasite absent** | **-0.86** | **0.20** | **-4.24** | **$2.2 \times 10^{-5}$** |
| Check 7 x parasite absent | -0.43 | 0.29 | -1.47 | 0.14 |
| **Check 7+ x parasite absent** | **-1.95** | **0.75** | **-2.59** | **$9.64 \times 10^{-3}$** |

| *Random effects* | *intercept variance* | *standard deviation* |
|---|---|---|
| well | 0.29 | 0.54 |

AIC: 1900.3

Table note: Pupation ~ checkpoint + parasite * heatwave + population + (heatwave * checkpoint) + (parasite * checkpoint) + (1 | well), modeled as a binomial distribution with log-log link using lme4.

**Table S31. Simplified pupation hazard robustness check emmeans pairwise contrasts for the population-level two-way interaction model (Table S30), with Bonferroni-adjusted p-values. P-values < 0.05 are indicated with asterisks.**

| Interaction | Populations | Checkpoint | Treatment | Estimate | SE | Z ratio | p-value |
|---|---|---|---|---|---|---|---|
| Checkpoint x heatwave | - | 4 | Early heatwave, control | 0.85 | 0.24 | 3.57 | $7 \times 10^{-4}$* |
| | - | 4 | Late heatwave, control | 0.81 | 0.24 | 3.3 | $1.7 \times 10^{-3}$* |
| | - | 5 | Early heatwave, control | 0.64 | 0.16 | 4.11 | $<1 \times 10^{-4}$* |
| | - | 5 | Late heatwave, control | 0.32 | 0.16 | 1.95 | 0.1 |
| | - | 6 | Early heatwave, control | 0.22 | 0.29 | 0.76 | 0.9 |
| | - | 6 | Late heatwave, control | -0.39 | 0.25 | -1.57 | 0.23 |
| | - | 7+ | Early heatwave, control | -1.35 | 1.15 | -1.17 | 0.48 |
| | - | 7+ | Late heatwave, control | -1.18 | 0.58 | -2.03 | 0.08 |
| Checkpoint x parasite exposure | - | 4 | Parasite present, absent | -1.16 | 0.19 | -6.09 | $<1 \times 10^{-4}$* |
| | - | 5 | Parasite present, absent | -0.29 | 0.13 | -2.25 | 0.02* |
| | - | 6 | Parasite present, absent | -0.72 | 0.24 | -2.99 | $2.8 \times 10^{-3}$* |
| | - | 7+ | Parasite present, absent | 0.79 | 0.73 | 1.08 | 0.28 |
| Population | Alameda, San Mateo | - | - | -0.54 | 0.17 | -3.14 | 0.01* |
| | Alameda, San Diego | - | - | 0.14 | 0.17 | 0.86 | 1.0 |
| | Alameda, Santa Barbara | - | - | -0.33 | 0.17 | -1.93 | 0.32 |
| | San Mateo, San Diego | - | - | 0.68 | 0.15 | 4.59 | $<1 \times 10^{-4}$* |
| | San Mateo, Santa Barbara | - | - | 0.2 | 0.15 | 1.32 | 1.0 |
| | San Diego, Santa Barbara | - | - | -0.48 | 0.15 | -3.18 | $9 \times 10^{-3}$* |

**Table S32. Host survival robustness check results estimating interactive effects of parasite exposure, heatwave treatment, and population on host survival to pupation. Significant predictors ($p < 0.05$) are highlighted with bold text.**

| *Predictors* | *Estimate* | *Standard Error* | *Z-statistic* | *p-value* |
|---|---|---|---|---|
| **(Intercept)** | **0.88** | **0.16** | **5.54** | **$3.08 \times 10^{-8}$** |
| **Early heatwave** | **1.42** | **0.32** | **4.39** | **$1.12 \times 10^{-5}$** |
| Late heatwave | 0.23 | 0.24 | 0.95 | 0.34 |
| **Parasite absent** | **2.45** | **0.50** | **4.88** | **$1.04 \times 10^{-6}$** |
| **Early heatwave x parasite absent** | **-1.50** | **0.71** | **-2.13** | **0.03** |
| Late heatwave x parasite absent | 0.28 | 1.20 | 0.24 | 0.81 |
| San Mateo | 0.40 | 0.28 | 1.44 | 0.15 |
| **San Diego** | **0.57** | **0.27** | **2.11** | **0.03** |
| Santa Barbara | 0.03 | 0.24 | 0.13 | 0.90 |
| Early heatwave x San Mateo | -0.19 | 0.52 | -0.36 | 0.72 |
| Late heatwave x San Mateo | -0.60 | 0.40 | -1.49 | 0.13 |
| **Early heatwave x San Diego** | **-1.12** | **0.49** | **-2.27** | **0.02** |
| Late heatwave x San Diego | -0.66 | 0.38 | -1.72 | 0.09 |
| Early heatwave x Santa Barbara | 1.24 | 0.68 | 1.82 | 0.07 |
| Late heatwave x Santa Barbara | 0.86 | 0.47 | 1.84 | 0.07 |
| Parasite absent x San Mateo | -0.22 | 0.80 | -0.28 | 0.78 |
| Parasite absent x San Diego | -1.33 | 0.67 | -1.99 | 0.05 |
| Parasite absent x Santa Barbara | 0.86 | 1.03 | 0.83 | 0.40 |
| Early heatwave x parasite absent x San Mateo | -0.25 | 1.11 | -0.23 | 0.82 |
| Late heatwave x parasite absent x San Mateo | -0.48 | 1.50 | -0.32 | 0.75 |
| Early heatwave x parasite absent x San Diego | 1.57 | 1.01 | 1.55 | 0.12 |
| Late heatwave x parasite absent x San Diego | 0.61 | 1.41 | 0.43 | 0.66 |
| Early heatwave x parasite absent x Santa Barbara | -1.55 | 1.43 | -1.08 | 0.28 |
| Late heatwave x parasite absent x Santa Barbara | -2.37 | 1.78 | -1.33 | 0.18 |

AIC: 266.16

Table note: Survival to pupation ~ heatwave treatment * parasite exposure * population, modeled as a binomial distribution using with weak priors on fixed effects using R package "arm".

**Table S33. Host survival robustness check emmeans pairwise contrasts for the population-level three-way interaction model (Table S32). Bonferroni-adjusted p-values <0.05 are marked with asterisks.**

| Predictors | Contrasts | | | Estimate | SE | Z ratio | p-value |
|---|---|---|---|---|---|---|---|
| | **Region or population** | **Heatwave treatment** | **Parasite** | | | | |
| Heatwave treatment x parasite exposure | - | control, early | present | 1.42 | 0.32 | 4.4 | $<1 \times 10^{-4}$* |
| | - | control | present, absent | -2.45 | 0.4 | -4.88 | $<1 \times 10^{-4}$* |
| | - | early | present, absent | -1.0 | 0.53 | 1.81 | 0.07 |
| Population x early heatwave | Alameda, San Mateo | early | - | -0.44 | 0.72 | -0.62 | 1.0 |
| | Alameda, San Diego | early | - | -0.04 | -0.7 | -0.05 | 1.0 |
| | Alameda, Santa Barbara | early | - | -1.39 | 0.89 | -1.57 | 0.7 |
| | San Diego, Santa Barbara | early | - | -1.35 | 0.8 | -1.68 | 0.55 |
| | San Diego, San Mateo | early | - | 0.41 | 0.61 | 0.66 | 1.0 |
| | Alameda | early, control | | 0.86 | 0.74 | 1.17 | 0.49 |
| | San Diego | early, control | - | 0.33 | 0.54 | 0.62 | 1.0 |
| Population x parasite exposure | Alameda, San Mateo | - | present | -0.99 | 0.31 | -3.23 | $7.4 \times 10^{-3}$* |
| | Alameda, San Diego | - | present | -0.83 | 0.29 | -2.88 | 0.02* |
| | Alameda, Santa Barbara | - | present | -1.58 | 0.39 | -4.02 | $3 \times 10^{-4}$* |
| | San Diego, San Mateo | - | present | 0.16 | 0.31 | 0.51 | 1.0 |
| | San Diego, Santa Barbara | - | present | -0.76 | 0.4 | -1.9 | 0.35 |
| | Alameda | - | present, absent | -3.57 | 1.59 | -2.25 | 0.02* |
| | San Mateo | - | present, absent | -1.58 | 0.51 | -3.1 | $1.9 \times 10^{-3}$* |
| | San Diego | - | present, absent | -1.44 | 0.44 | -3.25 | $1.2 \times 10^{-3}$* |
| | Santa Barbara | - | present, absent | -1.6 | 0.76 | -2.11 | 0.04* |

**Table S34. Simplified host survival robustness check results estimating interactive effects of parasite exposure and heatwave treatment, and additive effects of population on host survival to pupation. Significant predictors ($p < 0.05$) are highlighted with bold text.**

| *Predictors* | *Estimate* | *Standard Error* | *Z-statistic* | *p-value* |
|---|---|---|---|---|
| **(Intercept)** | **0.84** | **0.15** | **5.47** | **$4.62 \times 10^{-8}$** |
| **Early heatwave** | **1.36** | **0.29** | **4.67** | **$3.03 \times 10^{-6}$** |
| Late heatwave | 0.20 | 0.23 | 0.88 | 0.38 |
| **Parasite absent** | **2.44** | **0.48** | **5.05** | **$4.37 \times 10^{-7}$** |
| San Mateo | 0.15 | 0.17 | 0.89 | 0.37 |
| San Diego | 0.09 | 0.16 | 0.54 | 0.59 |
| **Santa Barbara** | **0.42** | **0.18** | **2.37** | **0.02** |
| **Early heatwave x parasite absent** | **-1.50** | **0.68** | **-2.19** | **0.03** |
| Late heatwave x parasite absent | -0.34 | 0.69 | -0.50 | 0.62 |

AIC: 410.51

Table note: Proportion larvae surviving to pupation ~ heatwave treatment * parasite exposure + population ID, modeled as a binomial distribution using base R function “glm()”.

**Table S35. Simplified host survival robustness check emmeans pairwise contrasts for the population-level two-way interaction model (Table S34). Bonferroni-adjusted p-values <0.05 are marked with asterisks.**

| | Contrasts | | | | | | |
|---|---|---|---|---|---|---|---|
| **Interaction** | **Population** | **Heatwave treatment** | **Parasite** | **Estimate** | **SE** | **Z ratio** | **p-value** |
| Heatwave treatment x parasite exposure | - | control, early | present | 1.36 | 0.29 | 4.67 | $<1 \times 10^{-4}$* |
| | - | control | present, absent | 2.44 | 0.48 | 5.05 | $<1 \times 10^{-4}$* |
| | - | early | present, absent | 0.94 | 0.49 | 1.95 | 0.05 |
| population | Alameda, San Mateo | - | - | -0.81 | 0.27 | -3.01 | 0.02* |
| | Alameda, San Diego | - | - | -0.74 | 0.26 | -2.88 | 0.02* |
| | Alameda, Santa Barbara | - | - | -1.08 | 0.28 | -3.83 | $8 \times 10^{-4}$* |
| | San Mateo, San Diego | - | - | 0.06 | 0.27 | 0.24 | 1.0 |
| | San Mateo, Santa Barbara | - | - | -0.27 | 0.29 | -0.94 | 1.0 |
| | San Diego, Santa Barbara | - | - | -0.33 | 0.28 | -1.21 | 1.0 |
